# Chemical Tools for the Gid4 Subunit of the Human E3 Ligase C-terminal to LisH (CTLH) Degradation Complex

**DOI:** 10.1101/2023.11.13.566858

**Authors:** Aliakbar Khalili Yazdi, Sumera Perveen, Xiaosheng Song, Aiping Dong, Magdalena M. Szewczyk, Matthew F. Calabrese, Agustin Casimiro-Garcia, Subramanyam Chakrapani, Matthew S. Dowling, Emel Ficici, Jisun Lee, Justin I. Montgomery, Thomas N. O’Connell, Grzegorz J. Skrzypek, Tuan P. Tran, Matthew D. Troutman, Feng Wang, Jennifer A. Young, Jinrong Min, Dalia Barsyte-Lovejoy, Peter J. Brown, Vijayaratnam Santhakumar, Cheryl H. Arrowsmith, Masoud Vedadi, Dafydd R. Owen

**Affiliations:** Structural Genomics Consortium, University of Toronto, Toronto, ON, Canada; Pfizer Research & Development, Groton, CT, USA; Pfizer Research & Development, Cambridge, MA, USA; Department of Pharmacology and Toxicology, University of Toronto, Toronto, ON, Canada; Princess Margaret Cancer Centre and Department of Medical Biophysics, University of Toronto, Toronto, ON, Canada

## Abstract

We have developed a novel chemical handle (PFI-E3H1) and a chemical probe (PFI-7) as ligands for the Gid4 subunit of the human E3 ligase CTLH degradation complex. Through an efficient initial hit-ID campaign, structure-based drug design (SBDD) and leveraging the sizeable Pfizer compound library, we identified a 500 nM ligand for this E3 ligase through file screening alone. Further exploration identified a vector that is tolerant to addition of a linker for future chimeric molecule design. The chemotype was subsequently optimized to sub-100 nM Gid4 binding affinity for a chemical probe. These novel tools, alongside the suitable negative control also identified, should enable the interrogation of this complex human E3 ligase macromolecular assembly.

## Introduction

Targeted protein degradation, whether exploiting proteolysis targeting chimeras (PROTACs) or molecular glues, is an increasingly common medicinal chemistry strategy for modulating protein function, including tackling so-called undruggable targets.^1^ Remarkably, the uptake and enthusiasm for these strategies remain predicated on established chemical equity for a very small number of the >600 human E3 ligases found within the gene family.^2^ Through a public-private partnership, we have undertaken an effort to identify druglike chemical matter for E3 ligases to enable the community as whole, particularly looking for alternatives to the established cereblon and VHL ligands that remain the workhorse proteins in the field.^3^ The collaboration was not structured to go beyond identifying novel E3 chemical matter or move on to prove any functional degradation possibilities through investigating chimeric molecules.

In this work we target Gid4, a sub-unit of the human CTLH complex which retains the GID-like ubiquitin ligase mechanism found in yeast.^4^ Much of the understanding of Gid4 comes from its role in yeast biology, and the behavior of its human ortholog found within the CTLH complex remains somewhat extrapolated from there.^5^ In response to changes in environmental glucose, yeast undergo a rapid switch from gluconeogenic to glycolytic processes.^6^ This metabolic change is enabled, in part, through targeted protein degradation of gluconeogenic enzymes conducted by the multi-subunit E3 ligase GID (Glucose-Induced Degradation) complex.^7, 8^ In line with other multi-subunit E3 ligases (ex: Cullin-RINGs), the GID complex likely represents a diverse set of assemblies built on a common scaffold.^9^ This is comprised of RING and RING-like subunits Gid2 and Gid9 in complex with core structural subunits Gid1, 5, and 8.^10, 11^ This assembly can then further enlist substrate recruiting domains including Gid4 and Gid10.^12^ One of the best characterized functions of the yeast GID complex is recognition of substrates containing an N-terminal proline degron through direct recruitment to the Gid4 subunit, including the metabolic enzyme Fructose-1,6-bisphosphatase 1 (Fbp1).^7^ Unexpectedly, this activity appears to require formation of higher-order GID supramolecular assemblies comprised of as many as twenty protein subunits. These higher order structures have been shown to enable multipronged targeting of oligomeric substrates (including Fbp1) providing an exciting clue into a deeper level of protein regulation by this macromolecular machine.^13^

The orthologous human complex, CTLH, retains a GID-like ubiquitin ligase mechanism, including the formation of supramolecular assemblies.^13^ In addition, the mammalian complex retains a structurally homologous Gid4 substrate receptor.^14^ In contrast to the yeast enzyme, however, the function of the human CTLH complex remains only partially understood. Deeper interrogation of both the native and non-native roles of the CTLH complex require novel tools in order to dissect function. The Gid4 substrate receptor represents a conserved and functionally relevant component of the GID/CTLH assembly. Previous work revealed that Gid4 contains a novel β-barrel fold comprised of eight closed antiparallel strands with an internal helical bundle insertion. Substrates bind within the central channel of this β-barrel such that they are positioned for ubiquitination by the active GID/CTLH complex.^14^ While metabolic regulation differs dramatically between yeast and mammals, the Gid4 substrate binding pocket remains highly conserved, providing an attractive ligandable site for the advancement of small molecule tools.

## Results and Discussion

In contrast to the NMR fragment and DNA-encoded library approaches employed by Gingras and Sicheri to identify a Gid4 binder with a *K*_d_ of 5.6 μM as determined by isothermal titration calorimetry,^15^ we conducted an unbiased hit ID campaign using affinity selection mass spectrometry (ASMS). Using 1300-fold compression and screening around 500,000 compounds in total, a small number of Gid4 binders were identified from the campaign. Given the knowledge of the N-end rule pathway responsible for recognizing substrates bearing N-terminal proline residues (Pro/N-degrons) in Gid4 from Min,^16^ truncated peptidic Gid4 degrons such as **1** have been identified. In light of the pharmacophore found within **1**, particularly the N-terminal proline, the significance of compound **2** as a hit from the ASMS campaign was striking. Although not an N-terminal proline, the stereochemically unspecified mixture of compound(s) **2** featured a benzylated N-terminal glycine as part of a Gly-Ala-Phe tripeptide. Based on compound **1** the predicted binding mode for compound **2** would require toleration of a benzyl group deep in the Gid4 substrate recognition pocket and the possibility for N-alkylated glycine to replace proline in N-terminal interactions. Compound **2** had been in the Pfizer compound library since 1993 and had been registered with no defined stereochemistry. The presumed mixture of compounds representing **2** was then screened in Gid4 fluorescence polarization (FP) and surface plasmon resonance (SPR) assays to confirm binding. The compound was a weak, but confirmed, binder in both assays (38 μM and 15μ M respectively). With three orthogonal methods confirming the hit now in place, rather than establish the exact stereochemical make up of **2** we proceeded to perform a simple substructure search for N-benzylated glycine amides within the full, multimillion Pfizer compound library, beyond the 500,000 molecules already screened in ASMS. This file screen was conducted at single point concentration in the FP assay of 50μM. Full *K* displacement (*K*_disp_) values were then determined for the most potent binders.

Amongst the hits identified was a further simplification of the pseudo-Ala-Phe segment of compound **2**. Compounds **3, 4** and **5** proved to be stronger binders, registering SPR binding *K*_D_ values in the 2.7-9 μM range. The very simple glycine-based pharmacophore, seen in file mining hits **3** and **4**, was well placed to move the lead matter away from the original tripeptide hit of compound **2** into more conventional, small molecule and druglike space.

Compound **4** with a *p*-methoxy benzyl (PMB) group on each end of a simple glycine amino acid core (and in the Pfizer file since 1987) was selected for crystallography. The co-crystal structure (Fig. 1, PDB:7S12) showed that the N-benzylated glycine did indeed perform the N-terminal proline role from the synthetic peptidic degrons already known (e.g. **1**). The basic amine showed a clear interaction with Glu-237 at the base of the degron pocket, designed to recognize an endogenous N-terminal proline. In now having an acyclic N-terminus, the presence of the N-benzylation was accommodated next to a wall of hydrophobic residues in the Gid-4 pocket (Leu-164, Leu-240 and Ile-249). In terms of the glycine component of the molecule, the amide carbonyl accepted a hydrogen bond from Gln-132 and donated an NH hydrogen bond to the backbone carbonyl of Ser-253 as it vectored the second PMB group out of the protein. This amidic PMB group did not display any obvious, significant Gid-4 interactions other than with Leu-164. This suggested the possibility of considerable flexibility in other groups that could play the role of the exit vector and reinforced the efficiency of the proline mimetic interactions deeper in the degron N-terminus pocket.

**Figure 1.**
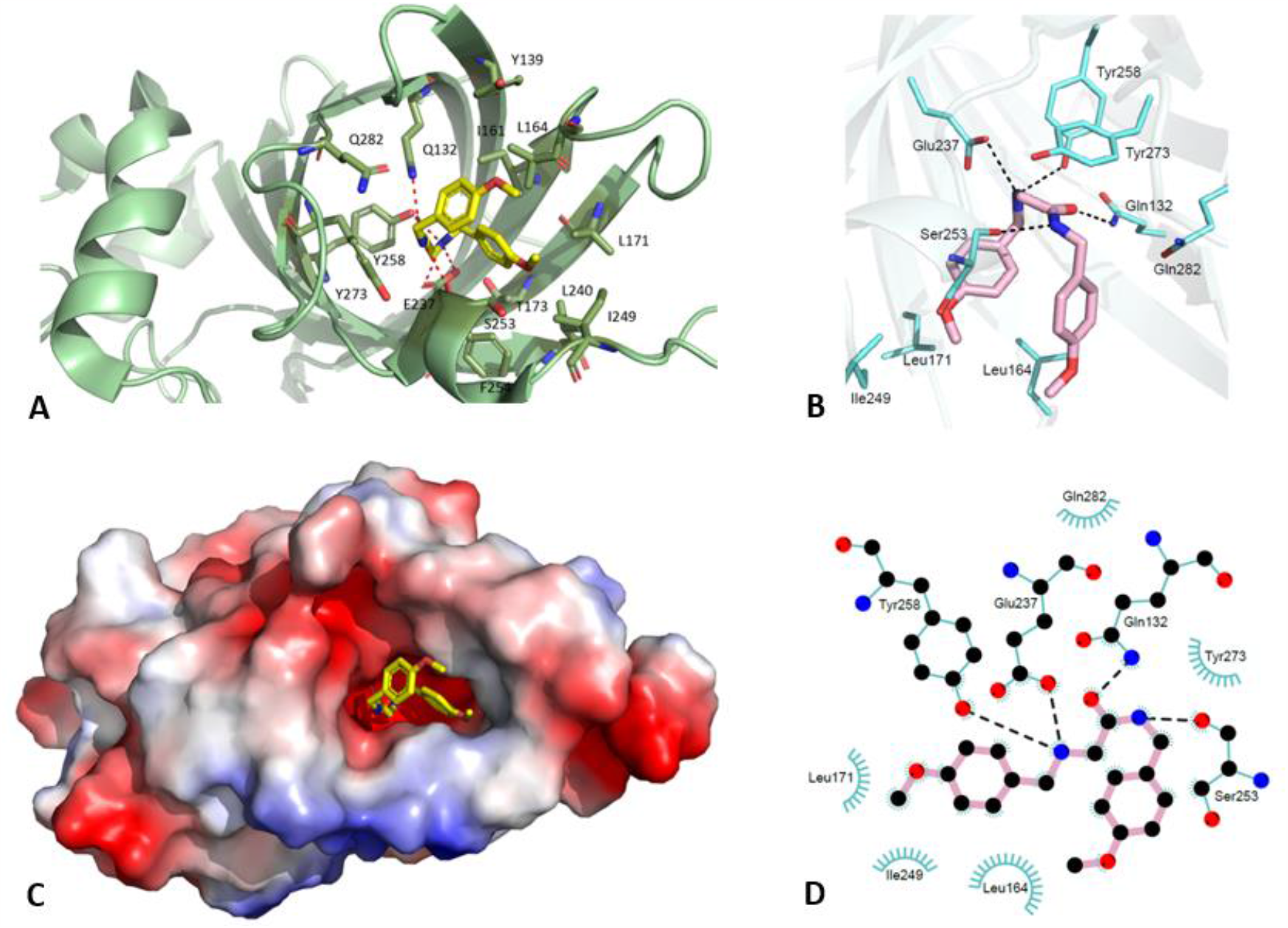
Structural biology for compound 4: **A** View from exit vector for *p*-methoxy benzylamide substituent; **B** Critical polar interactions in degron N-terminal proline binding site, in particular N-benzylglycine substituent of compound **4**; **C** Protein space filling model viewed from ligand exit vector; **D** Two-dimensional cartoon view of key polar/non-polar protein interactions for compound **4** (pink)

As benzylamides of glycine were established as tolerated, a file screen of available proline benzylamides was conducted. While compounds such as **6** do bind, it appears there is an advantage to using the benzylated and acyclic glycine as a proline mimetic to capitalize on the leucine-rich wall in the pocket. One further sub-structural file screen was carried out where N-benzylated glycines with a diversity of more complex exit vectors were tested (beyond just the simple benzylamide such as in **3**). This identified a number of compounds such as **7** which all contained a characteristic *cis*-1,4-substituted cyclohexane. Compound **7** had a *K*_D_ of 0.5 μM in SPR binding to Gid4. It is worth reflecting that no compounds had been designed or synthesized up to this point in the project. The SAR learnings were entirely derived from compounds retained in the corporate library collection, some of which had been first registered at Pfizer as long ago as 1958 (e.g. **3**). Compound **7** (now renamed PFI-E3H1) met our agreed collaboration endpoint of being an ‘E3 handle’. We had set out to identify potent (∼500 nM), tractable chemical matter displaying tangible SAR, with the potential for cell penetration for a novel E3 ligase. PFI-E3H1 (**7**) met these criteria.

In order to confirm cellular penetration and target engagement for the chemotype, we performed a NanoLuciferase (NanoLuc) bioluminescence resonance energy transfer (NanoBRET) assay, which measures protein-protein interaction in cells.^17^ NanoLuc-tagged with MPGLWKS degron (energy donor) and HaloTag-tagged Gid4, which binds to cell-permeable chloroalkane-modified 618 nm fluorophore energy acceptor, were expressed in HEK293T cells (Fig. 2A). The protein interaction is measured by determining NanoBRET ratio, calculated by dividing the acceptor emission value by the donor emission value. Cell treatment with compound **7** resulted in a dose-dependent decrease in Gid4 binding to MPGLKWS peptide with IC_50_ of 2.5 μM (Fig. 2B), indicating no significant barrier to cellular penetration with respect to its recombinant protein Gid4 SPR binding *K*_D_ determination of 0.5 μM.

**Figure 2.**
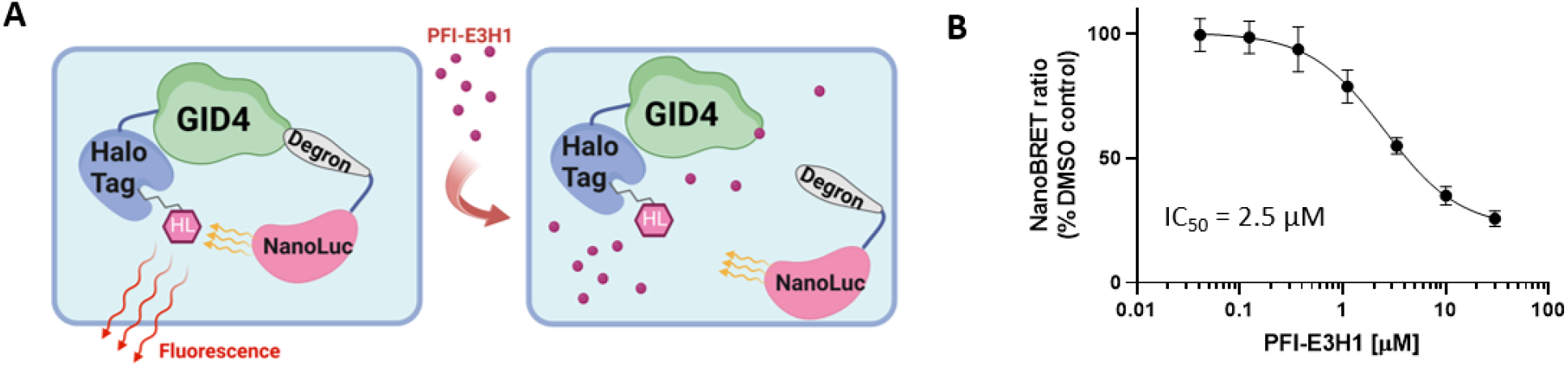
PFI-E3H1 (7) NanoBRET assay measuring the Gid4/MPGLWKS interaction: **A** Schematic representation of GID4 NanoBRET assay; **B** HEK293T cells were transfected with C-terminally NanoLuc-tagged MPGLWKS and N-terminally HaloTag-tagged GID4 for 24 h and treated with compound for 4 h. The graph (mean ± SD) represents non-linear fit of NanoBRET ratio normalized to DMSO control: *n* = 12, three separate experiments, IC_50_ = 2.5 ± 0.4 μM.

Based on the crystal structure of compound **4**, we synthesized one compound to emphasize a tolerated exit vector for those interested in the potential for chimeric molecules derived from the chemotype. As designed, compound **8** tolerated the presence of a PEG chain off the benzimidazole at the 5-position. With an E3 handle and a validated exit vector identified to enable investigation of any Gid4-related PROTAC hypotheses (Compounds **7** and **8**), we then moved onto designing a chemical probe for Gid4 itself. A cell penetrant, sub-100nM chemical probe enabled by structural biology would likely enhance the understanding of Gid4 biology, particularly around endogenous binding proteins and whether Gid4 plays any degradative or non-degradative functions. The simplicity of the chemotype made for a synthetic strategy focused on exploring the full potential of the N-terminal binding site, through reacting a number of benzyl primary amines with an α-chloroamide template. This was sufficiently robust to run in a library chemistry format.

**Scheme 1.**
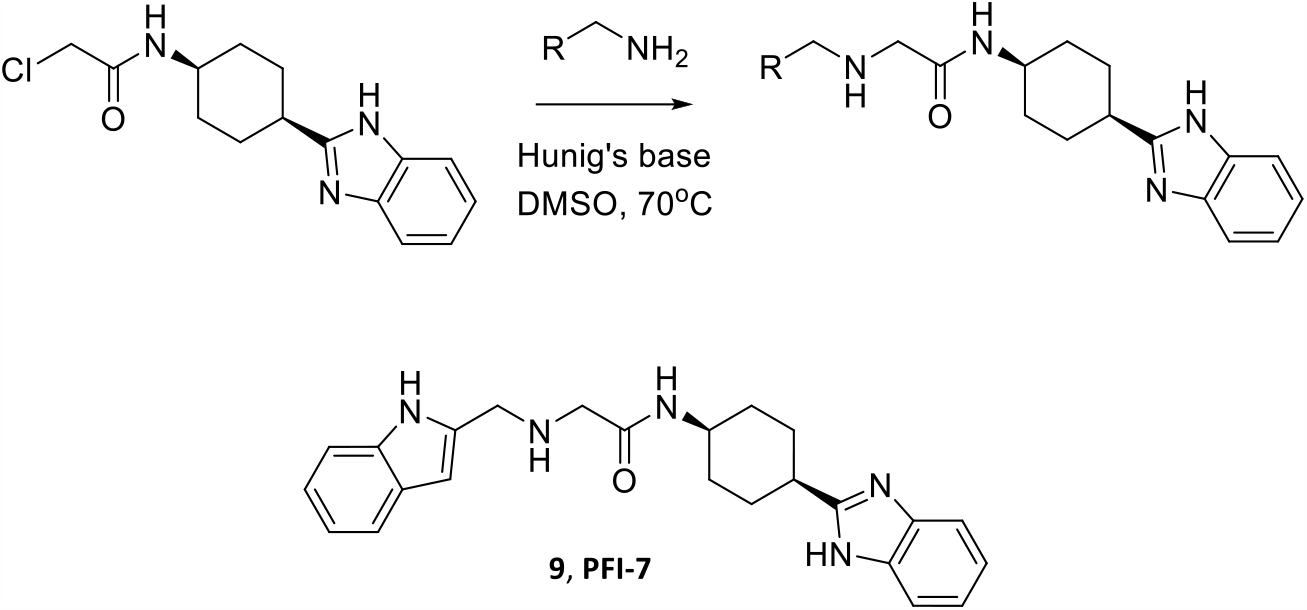
Library synthesis enabling SAR exploration of degron N-terminal mimetic

With a library success rate of 74%, 91 analogues were synthesized and tested where we were seeking the most potent binder by SPR that still retained physicochemical properties compatible with cellular penetration. Compound **9**, synthesized via library chemistry (Scheme 1), was an almost 6x improved binder compared to **7** by SPR (Table 1). Renamed as PFI-7, this molecule meets the commonly adopted criteria for a chemical probe.^18^ An important complement to any chemical probe is a suitable, structurally related negative control. Given the importance of the Glu-237 interaction with a secondary amine within the small molecule binding pharmacophore, capping this amine as an acetyl amide rendered compound **10** inactive against Gid4 (Table 1). This was then designated as the negative control (PFI-7N) for use in conjunction with the Gid4 chemical probe PFI-7.

**Table 1.**
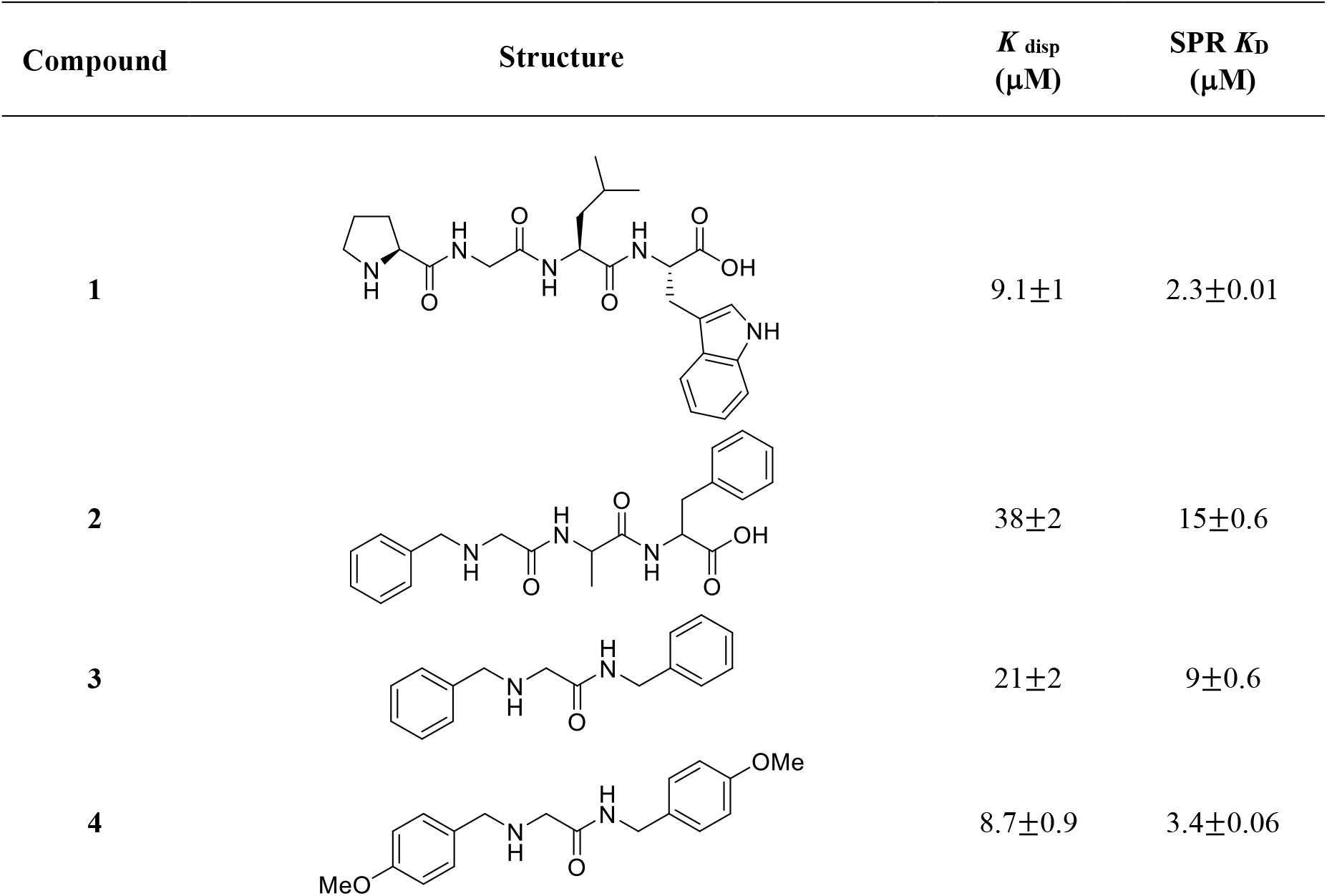
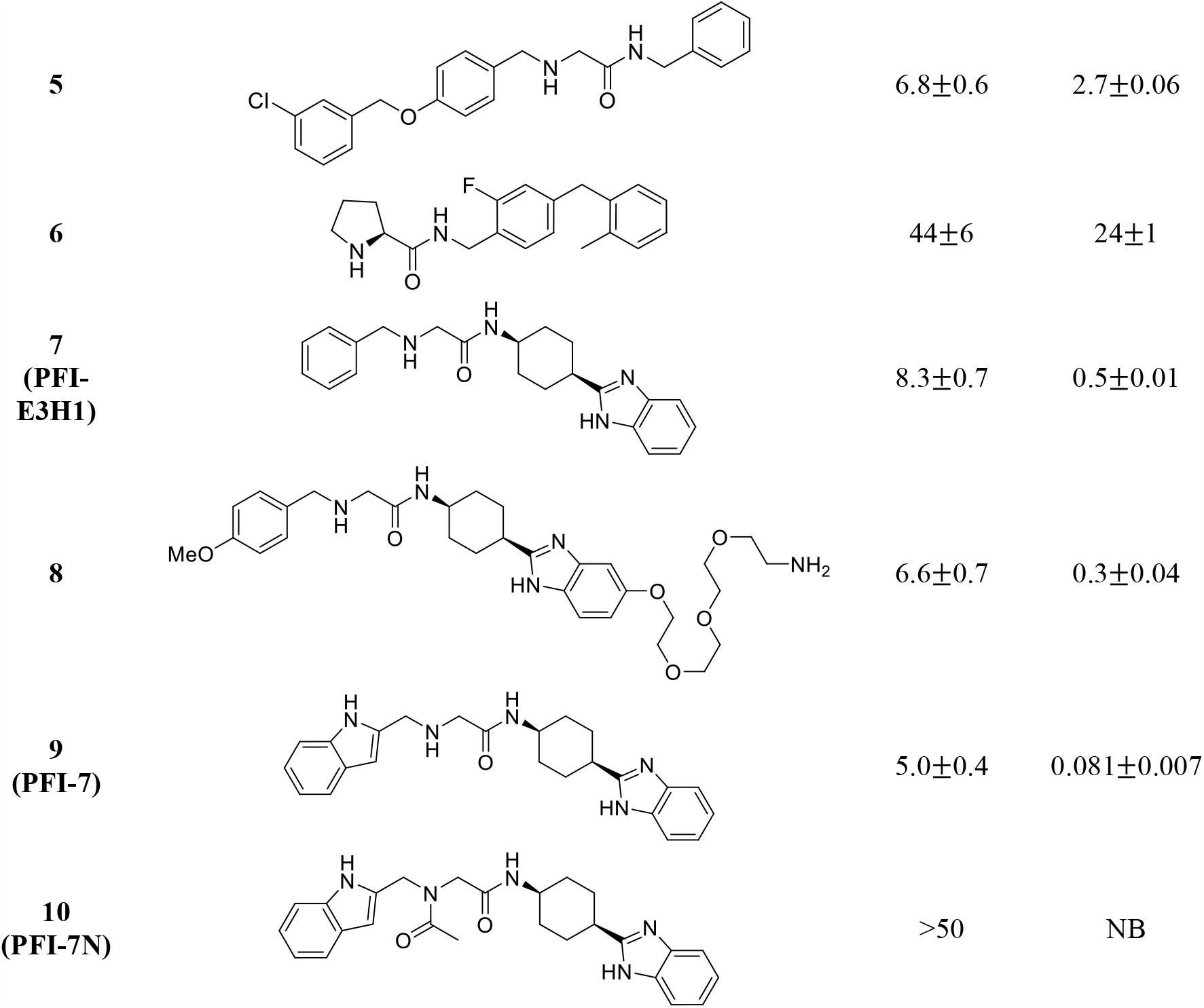
Gid4 binding characterization of compounds. *K*_disp_ values were determined using FP-based peptide displacement assay and *K*_D_ values using SPR steady state affinity fitting. All values represent the mean of three independent experiments ± standard deviation. NB: No Binding

| Compound | Structure | $K_{\text{disp}}$<br>(μM) | SPR $K_D$<br>(μM) |
| --- | --- | --- | --- |
| **1** | | 9.1±1 | 2.3±0.01 |
| **2** | | 38±2 | 15±0.6 |
| **3** | | 21±2 | 9±0.6 |
| **4** | | 8.7±0.9 | 3.4±0.06 |
| 5 |  | 6.8±0.6 | 2.7±0.06 |
| 6 |  | 44±6 | 24±1 |
| 7<br>(PFI-E3H1) |  | 8.3±0.7 | 0.5±0.01 |
| 8 |  | 6.6±0.7 | 0.3±0.04 |
| 9<br>(PFI-7) |  | 5.0±0.4 | 0.081±0.007 |
| 10<br>(PFI-7N) |  | >50 | NB |

## Conclusion

In summary, we have identified and optimized novel chemical matter for the Gid4 subunit of the human E3 ligase CTLH degradation complex through an efficient hit ID and optimization strategy. This chemotype represents an addition to the relatively small number of non-covalent E3 ligase ligands previously known. The molecules are of sufficient potency and physicochemical properties to yield tools (E3 Handle, PFI-E3H1, **7**) and a chemical probe (PFI-7, **9**) that will assist in the exploration of Gid4 biology.^19^

## Supporting information

GID Data and SI

## Author Contributions

**Program conception:** Matthew F. Calabrese, Dafydd R. Owen, Cheryl H. Arrowsmith

**Chemistry, Computational and Protein Science**: Agustin Casimiro-Garcia, Subramanyam Chakrapani, Matthew S. Dowling, Emel Ficici, Jisun Lee, Justin I. Montgomery, Thomas N. O’Connell, Grzegorz J. Skrzypek, Tuan P. Tran, Matthew D. Troutman, Feng Wang, Jennifer A. Young

**Crystallography**: Xiaosheng Song, Aiping Dong

**Biophysical assays**: Aliakbar Kahlili Yazdi, Sumera Perveen

**Cell biology:** Magdalena M. Szewczyk

**SGC University of Toronto PIs**: Cheryl H. Arrowsmith, Dalia Barsyte-Lovejoy, Peter J. Brown, Jinrong Min, Vijayaratnam Santhakumar, Masoud Vedadi

**Paper writing and figures:** Dalia Barsyte-Lovejoy, Matthew F. Calabrese, Dafydd Owen, Sumera Perveen, Jisun Lee, Magdalena M. Szewczyk

## Acknowledgements

The Structural Genomics Consortium is a registered charity (no: 1097737) that receives funds from Bayer AG, Boehringer Ingelheim, Bristol Myers Squibb, Genentech, Genome Canada through Ontario Genomics Institute [OGI-196], EU/EFPIA/OICR/McGill/KTH/Diamond Innovative Medicines Initiative 2 Joint Undertaking [EUbOPEN grant 875510], Janssen, Merck KGaA (aka EMD in Canada and US), Pfizer and Takeda.

