## Supplementary material for "Chemical Tools for the Gid4 Subunit of the Human E3 Ligase C-terminal to LisH (CTLH) Degradation Complex": GID Data and SI

Experimental Procedures and Supplementary Information

### Materials and Methods

All chemicals, reagents and solvents were purchased from commercial sources when available and used without further purification. Concentration *in vacuo* means that a rotary evaporator was used.

Proton (1H NMR) and carbon ($^{13}C$ NMR) nuclear magnetic spectroscopy were recorded with 400 MHz Bruker or Bruker Avance NEO 500 MHz Spectrometer with 5 mm DCH CryoProbe ($^{13}C$ observe probe) spectrometers. Chemical shifts are expressed in parts per million downfield with respect to solvent resonance as the internal standard (for example, $CDCl_3$ at 7.26 ppm for 1H and $CDCl_3$ at 77.16 ppm for 13C). The peak shapes are denoted as follows: s, singlet; d, doublet; t, triplet; q, quartet; p, pentet; m, multiplet; br s, broad singlet; dd, doublet of doublets, dt, doublet of triplets; dq, doublet of quartets. Silica gel chromatography was performed using a medium pressure ISCO system using columns pre-packaged by ISCO. Liquid chromatography mass spectrometry (LCMS) was performed on a Waters Acquity SQ with ZSpray dual ESI/APCI source (Waters Xselect HSS TS column, 30 x 2.1 mm, 2.5 µm particles or Waters Acquity BEH C18 column, 50 x 2.1 mm, 2.5 µm particles; 95% water/acetonitrile linear gradient to 5% water/acetonitrile over 0.7 min, hold at % water/acetonitrile to 1.1 min, 0.1% formic acid or 0.1% ammonium hydroxide modifier; flow rate of 0.8 mL/ min; 30°C). Mass spectrometry data are reported from LCMS analyses. Mass spectrometry (MS) was performed via atmospheric pressure chemical ionization (APCI), electrospray Ionization (ESI), electron impact ionization (EI) or electron scatter (ES) ionization sources. For high resolution mass spectrometry (HRMS), all data were gathered on a Sciex TripleTOF 5600+ (Sciex, Ontario, Canada) with DuoSpray ionization source. The LC instrument includes an Agilent (Agilent Technologies, Wilmington, DE) 1200 binary pump, Agilent 1200 autosampler, Agilent 1200 column compartment, and Agilent 1200 DAD. The instrument acquisition and data handling were done with Sciex Analyst TF version 1.7.1. Prior to acquisition instrument was calibrated with less than 5 ppm accuracy. During acquisition, a calibration run was performed initially and after every 5 injections using the Sciex positive polarity tuning mix. Elution Conditions: Column: Waters XSelect HSS T3, 2.1x50mm,

2.5µm particle size; Column Temperature 60 °C, Solvent A: Water (0.1% formic acid) , Solvent B: Acetonitrile (0.1% formic acid); Gradient: Initial 5% B, hold for 0.10 min, 5-95% B in 2.8 min, , 95-5% B in 0.20 min, 3.5 min total runtime; Flow rate 0.8 mL/min ; TOF Conditions: ESI in Positive Mode; The spray chamber: Gas 1 and 2 at 60, curtain gas at 40, temperature 600°C, IonSpray voltage 5500 V, declustering potential 100, collision energy 10. The acquisition is done in TOF MS mode with range of 100-2000 amu with accumulation time of 0.20 secs. ESI in Negative Mode; The spray chamber: Gas 1 and 2 at 60, curtain gas at 40, temperature 600°C, IonSpray voltage -4500 V, declustering potential -100, collision energy -10. The acquisition is done in TOF MS mode with range of 100-2000 amu with accumulation time of 0.20 secs.

**Preparation of compounds:**

**Compound 1:** CAS No. 2480140-84-7

**Compound 2:** CAS No.  97657-94-8

**Compound 3:** CAS No.  1089-31-2

**Compound 4:** CAS No.  117471-50-8

**Compound 5:** N-benzyl-2-((4-((3-chlorobenzyl)oxy)benzyl)amino)acetamide

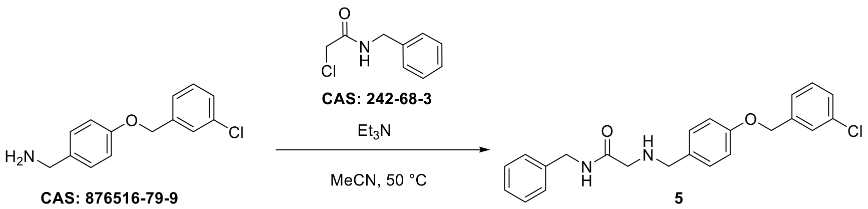


To a solution of (4-((3-chlorobenzyl)oxy)phenyl)methanamine (50 mg, 0.18 mmol) and triethylamine (74 µL, 0.528 mmol) in anhydrous acetonitrile (1.2 mL) was added 2-chloron-benzylacetamide (39 mg, 0.211 mmol). The resulting mixture was brought to 50 °C and left to stir overnight until conversion was complete. The reaction mixture was cooled down to room temperature and concentrated *in vacuo*. Purification by flash column chromatography on silica with 0 → 5% MeOH/DCM afforded **5** as a yellow oil (39 mg, 55%).

$^{1}H$ NMR (600 MHz, DMSO) δ 9.51 (s, 1H), 9.07 (t, *J* = 5.9 Hz, 1H), 7.52 (s, 1H), 7.46 (d, *J* = 8.3 Hz, 2H), 7.44 – 7.38 (m, 3H), 7.36 – 7.30 (m, 2H), 7.30 – 7.23 (m, 3H), 7.05 (d, *J* = 8.3 Hz, 2H), 5.15 (s, 2H), 4.33 (d, *J* = 5.9 Hz, 2H), 4.10 (s, 2H), 3.68 (s, 2H), 3.36 (s, residual $H_2O$). $^{13}C$ NMR (151 MHz, DMSO) δ 164.89, 158.51, 139.49, 138.51, 133.11, 131.85, 130.39, 128.30, 127.78, 127.37, 127.27, 126.99, 126.16, 123.83, 114.84, 68.24, 49.25, 46.40, 42.17. HRMS (ESI): calculated for ($C_{23}H_{23}ClN_2O_2$) $[M+H]^+$: 395.1521, found: 395.1527.

**Compound 6:** (S)-N-(2-fluoro-4-(2-methylbenzyl)benzyl)pyrrolidine-2-carboxamide

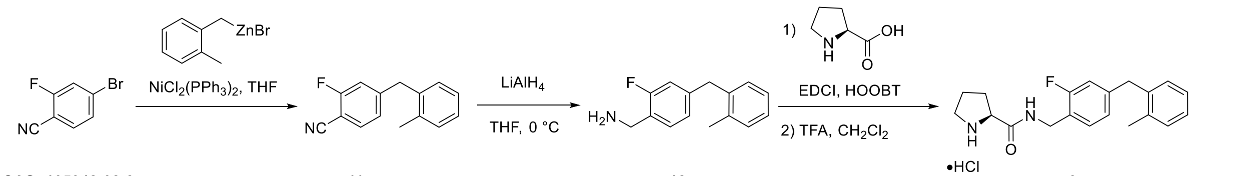

**CAS: 105942-08-3** **11** **12** **6**

Step 1: To a flame-dried flask under nitrogen charged with 4-bromo-2-fluorobenzonitrile (1.0 g, 5 mmol) in anhydrous THF (5 mL) was added 2-methylbenzyl zinc bromide (0.5 M in THF, 14 mL, 7 mmol), followed by $NiCl_2(PPh_3)_2$ (65 mg, 0.1 mmol). The resulting mixture was stirred under nitrogen for 1 h until conversion was complete. The reaction mixture was poured into 1 N HCl (20 mL), diluted with diethyl ether (30 mL) and the resulting layers were separated. The organic layer was then washed with water (20 mL), saturated $NaHCO_3$ (20 mL), brine (20 mL), and dried over $Na_2SO_4$, filtered, and concentrated *in vacuo* to afford **11** (1.46 g) which was used in the subsequent step without further purification.

Step 2: To a solution of lithium aluminum hydride in THF (1 M in THF, 15 mL, 15 mmol) at 0 °C was added dropwise solution of **11** (1.46 g) in anhydrous THF (5 mL). The resulting mixture was stirred at 0 °C for 1 h until conversion was complete. The reaction flask was then open to air and quenched with slow addition of water (0. 57 mL) at 0 °C before adding 15% NaOH (aq., 0.57

mL). The reaction mixture was further diluted with 1:1 water/THF (4 mL), filtered, and concentrated *in vacuo* to afford **12** (1.38 g) that was used without further purification.

Step 3: EDCI (1.44 g, 7.5 mmol) and HOOBt (1.22 g, 7.5 mmol) were added to a solution of *N*-Boc-L-proline (1.61 g, 7.5 mmol) in $CH_2Cl_2$ (7.5 mL) and DMF (7.5 mL) at 0 °C. The resulting mixture was left to stir at 0 °C for 30 min before **12** (1.38 g) was added as a solution in DMF (2 mL). The reaction mixture was then warmed to room temperature and stirred until complete conversion of the amine was observed by LCMS. The mixture was poured into water (15 mL) and extracted with diethyl ether (30 mL, x2). The organic layers were combined and washed with sat. $NaHCO_3$ (20 mL), brine (20 mL), and dried over $Na_2SO_4$, filtered, and concentrated *in vacuo*. The crude material was immediately taken up in a 1:1 mixture of $CH_2Cl_2$/TFA (10 mL) and stirred at room temperature for 30 min. The reaction mixture was then concentrated *in vacuo*, diluted with 1N NaOH (aq., 10 mL) and extracted with diethyl ether (20 mL). The organic layer was dried over $Na_2SO_4$, filtered, and concentrated *in vacuo*. Purification by flash column chromatography on silica with 45:45:10 EtOAc/$CH_2Cl_2$/MeOH afforded the desired product as the free base which was then converted to its HCl salt and triturated with EtOAc to give **6** as a white solid (0.66 g, 36%). $^1$H NMR (600 MHz, DMSO) δ 9.19 (t, *J* = 5.7 Hz, 1H), 7.27 (t, *J* = 7.9 Hz, 1H), 7.21 – 7.07 (m, 5H), 6.97 – 6.90 (m, 2H), 4.38 – 4.27 (m, 2H), 4.19 (t, *J* = 7.7 Hz, 1H), 3.95 (s, 2H), 3.36 (s, residual $H_2O$), 3.24 – 3.13 (m, 2H), 2.34 – 2.26 (m, 1H), 2.19 (s, 3H), 1.92 – 1.78 (m, 3H). $^{13}$C NMR (151 MHz, DMSO) δ 168.10, 160.82, 159.19, 142.32, 142.27, 138.36, 136.03, 136.01, 135.95, 130.17, 129.71, 129.68, 129.66, 126.51, 126.02, 124.51, 124.49, 122.55, 122.45, 115.18, 115.04, 58.75, 45.37, 37.86, 36.18, 36.15, 29.64, 23.57, 19.19. HRMS (ESI): calculated for ($C_{20}H_{23}FN_2O$) $[M+H]^+$: 327.1867, found: 327.1859.

**Compound 7: PFI-E3H1**; N-(*cis*-(1,4)-4-(1H-benzo[d]imidazol-2-yl)cyclohexyl)-2-(benzylamino)acetamide

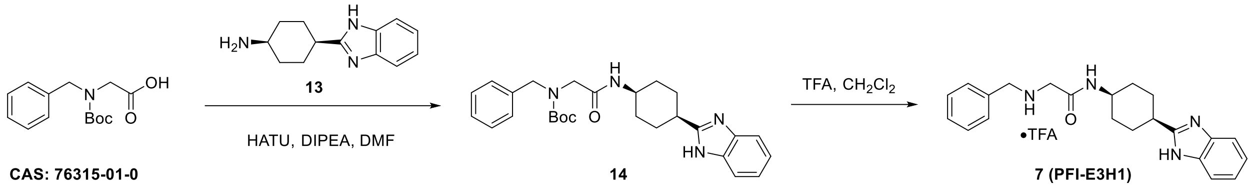


To a mixture of N-benzyl-N-(tert-butoxycarbonyl)glycine (200 mg, 0.754 mmol), **13** (239 mg, 0.829 mmol), and DIPEA (0.53 mL, 3.02 mmol) in anhydrous DMF (1.51 mL) was added HATU (344 mg, 0.905 mmol). The resulting mixture was left to stir at room temperature overnight until conversion was complete. The reaction mixture was then poured into 1N HCl (5 mL) and extracted with $CH_2Cl_2$ (10 mL, x2) before combining the organic layers. Dried the organic layer with sat. $NaHCO_3$ (5 mL) and dried over $Na_2SO_4$, filtered, and concentrated *in vacuo*. Purification by flash column chromatography on silica with 4:1 → 0.1 n-heptanes/EtOAc afforded **14** as a white solid which was immediately used in the subsequent step by dissolving the solid in $CH_2Cl_2$ (3 mL) and TFA (0.3 mL, 3.77 mmol). The resulting mixture was left to stir at room temperature for 17h. The reaction mixture was evaporated under reduced pressure and dried under high-vacuum to give **7** as a TFA salt (349 mg, 78%). $^1$H NMR (500 MHz, DMSO) δ 12.09 (s, 1H), 7.79 – 7.66 (m, 1H), 7.54 – 7.36 (m, 2H), 7.31 – 7.18 (m, 4H), 7.18 – 7.12 (m, 1H), 7.12 – 7.04 (m, 2H), 3.86 – 3.78 (m, 1H), 3.62 (s, 2H), 3.14 (s, 1H), 3.03 (s, 2H), 3.00 – 2.92 (m, 1H), 2.10 – 2.00 (m, 2H), 1.87 – 1.77 (m, 2H), 1.66 – 1.54 (m, 4H). $^{13}$C NMR (126 MHz, DMSO) δ 169.99, 169.97, 157.80, 140.12, 128.20, 128.13, 128.11, 126.76, 121.11, 52.64, 51.34, 48.64, 44.87, 40.43, 34.82, 28.99, 26.44. HRMS (ESI): calculated for ($C_{22}H_{26}N_4O$) $[M+H]^+$: 363.2179, found: 363.2171.

**Compound 13:** Synthesis of *cis*-(1,4)-4-(1H-benzo[d]imidazol-2-yl)cyclohexan-1-amine •2HCl

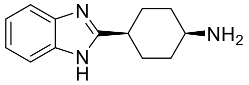

To a solution of (1s,4s)-4-aminocyclohexane-1-carboxylic acid (265 g, 1.85 mol) in 6N aq. HCl (5.3 L) was added 1,2-diaminobenzene (204 g, 1.89 mol) at room temperature. The resulting mixture was heated to reflux overnight until TLC (Ethyl acetate/Petroleum ether, 1:4) indicated

the reaction was complete. The reaction mixture was concentrated *in vacuo* to give 350 g of crude product which was washed with a mixture of methanol (500 mL) and water (30 mL) twice to afford **13** as a pink solid (180 g, 34%). LCMS m/z 216 $[M+H]^+$. $^1$H NMR (400 MHz, Methanol-$d_4$) δ 7.89 – 7.72 (m, 2H), 7.62 (dt, *J* = 6.2, 3.4 Hz, 2H), 3.59 (tt, *J* = 7.1, 4.3 Hz, 1H), 3.50 (tt, *J* = 7.6, 4.1 Hz, 1H), 2.42 (dtd, *J* = 14.9, 7.5, 3.7 Hz, 2H), 2.22 (ddt, *J* = 13.5, 8.5, 4.2 Hz, 2H), 2.11 (ddt, *J* = 12.6, 8.2, 4.1 Hz, 2H), 1.86-1.66 (m, 2H).

**Compound 8:** N-((1s,4s)-4-(6-(2-(2-(2-(2-aminoethoxy)ethoxy)ethoxy)ethoxy)-1H-benzo[d]imidazol-2-yl)cyclohexyl)-2-((4-methoxybenzyl)amino)acetamide hydrochloride salt

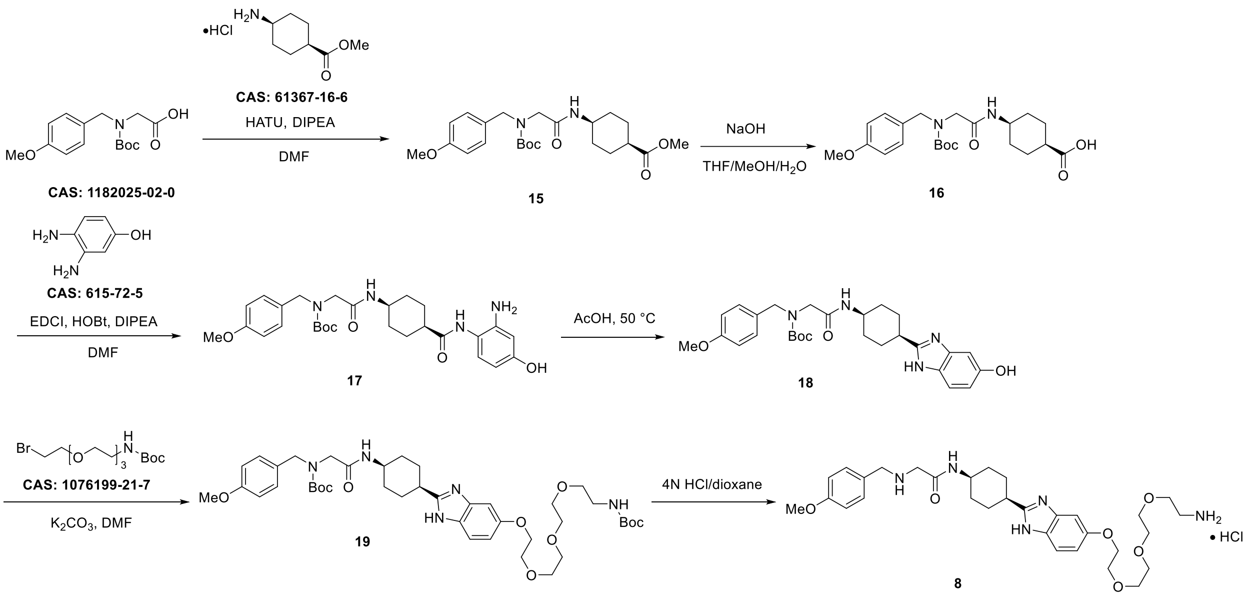


Step 1: A solution of *N*-(tert-butoxycarbonyl)-N-(4-methoxybenzyl)glycine (100 mg, 0.339 mmol) and HATU (193 mg, 0.508 mmol) in anhydrous DMF (3 mL) was stirred at room temperature for 15 min before methyl (1s,4s)-4-aminocyclohexane-1-carboxylate hydrochloric acid salt (86 mg, 0.372 mmol) and hunig's base (0.173 mL, 1.02 mmol) was added. The resulting mixture was left to stir at room temperature for 16 h. The reaction mixture was diluted with water (15 mL) and extracted with EtOAc (10 mL x 3). The organic layers were then combined and washed with brine, dried over $Na_2SO_4$, filtered, and concentrated *in vacuo*.

Purification by flash column chromatography on silica gel with 0 → 20% EtOAc/n-heptanes gave **15** as a colorless oil (100 mg, 68%). $^{1}$H NMR (400 MHz, MeOD) δ 7.98 (s, residual DMF), 7.19 (d, *J* = 8.7 Hz, 2H), 6.89 (d, *J* = 8.7 Hz, 2H), 5.49 (s, residual DCM), 4.87 (s, residual $H_2O$), 4.43 (s, 2H), 3.78 (m, 6H), 3.67 (s, 3H), 3.00 (s, residual DMF), 2.86 (s, residual DMF), 2.56 – 2.48 (m, 1H), 1.97 – 1.81 (m, 2H), 1.72 – 1.61 (m, 4H), 1.52 – 1.44 (m, 11H).

Step 2: To a solution of **15** (100 mg, 0.23 mmol) in THF (1 mL), MeOH (1 mL), and water (0.25 mL) was added solid NaOH (18.4 mg, 0.46 mmol). The resulting mixture was left to stir at room temperature for 1 h in which TLC (1:1 EtOAc/$CH_2Cl_2$) analysis against the starting material showed complete consumption of the starting material. The reaction mixture was then diluted with water (2 mL) and the pH was adjusted to 3 using aqueous 10% citric acid. The aqueous solution was extracted with EtOAc (15 mL x 3) before combining the organic layers which was then washed with brine, dried over $Na_2SO_4$, filtered, and concentrated *in vacuo* to afford **16** as a colorless gum (97 mg, quantitative). $^{1}$H NMR (400 MHz, MeOD) δ 7.19 (d, *J* = 8.6 Hz, 2H), 6.89 (d, *J* = 8.7 Hz, 2H), 4.88 (s, residual $H_2O$), 4.43 (s, 2H), 4.10 (q, J = 7.2, residual EtOAc), 3.84 – 3.67 (m, 6H), 2.52 – 2.43 (m, 1H), 2.01 (s, residual EtOAc), 1.97 – 1.82 (m, 2H), 1.71 – 1.61 (m, 4H), 1.56 – 1.43 (m, 11H), 1.24 (t, J = 7.1, residual EtOAc).

Step 3: A mixture of **16** (97 mg, 0.23 mmol), HOBt (46.8 mg, 0.35 mmol), and EDCI (66.3 mg, 0.35 mmol) in anhydrous DMF (2 mL) was stirred at room temperature for 15 min before adding 3,4-diaminophenol (102 mg, 0.58 mmol) and DIPEA (89.4 mg, 0.69 mmol) dropwise. The resulting brown mixture was left to stir at room temperature for 16 h. The reaction mixture was diluted with water (5 mL) and extracted with EtOAc (15 mL x 3). The organic layers were combined and washed with brine, dried over $Na_2SO_4$, filtered, and concentrated *in vacuo*. The crude product was then purified using flash column chromatography on silica gel with 0 → 10% MeOH/$CH_2Cl_2$ to afford **17** as a brown solid (60 mg, 49%). $^{1}$H NMR (400 MHz, $CDCl_3$) δ 7.24 – 7.14 (m, 2H), 6.91 – 6.81 (m, 3H), 6.29 – 6.20 (m, 2H), 5.30 (s, residual DCM), 4.42 (s,

2H), 3.99 – 3.89 (m, 1H), 3.82 – 3.73 (m, 5H), 2.96 (s, residual DMF), 2.88 (s, residual DMF), 2.42 – 2.31 (m, 1H), 1.76 – 1.54 (m, 8H), 1.48 (s, 9H).

Step 4: A solution of **17** (60 mg, 0.11 mmol) in acetic acid (0.65 mL) was stirred at 50 °C for 32 h. The reaction mixture was concentrated *in vacuo* to afford the crude product, which was then dissolved in 1:1 EtOAc/aq. $NaHCO_3$ (2 mL) and was stirred at room temperature for ~10 min. The mixture was diluted with EtOAc (2 mL) before separating the layers. The aqueous layer was extracted with EtOAc (2 mL x 3), and the combined organic layers were washed with brine, dried over $Na_2SO_4$, filtered, and concentrated to afford **18** (56 mg, quantitative) which was used in the subsequent step without further purification. $^1$H NMR (400 MHz, MeOD) δ 7.58 (s, 1H), 7.42 (d, *J* = 8.8 Hz, 1H), 7.21 – 7.13 (m, 2H), 6.97 (d, *J* = 2.2 Hz, 1H), 6.89 – 6.82 (m, 3H), 4.89 (s, residual $H_2O$), 4.42 (s, 2H), 4.10 (q, J = 7.2, residual EtOAc), 4.00 – 3.94 (m, 1H), 3.87 – 3.68 (m, 5H), 3.22 – 3.09 (m, 1H), 2.16 – 2.00 (m, 2H), 2.01 (s, residual EtOAc), 1.99 (s, residual AcOH), 1.86 – 1.62 (m, 5H), 1.54 – 1.37 (m, 10H), 1.24 (t, J = 7.1, residual EtOAc).

Step 5: To a solution of **18** (56 mg, 0.11 mmol) in anhydrous DMF (0.31 mL) was added potassium carbonate (46 mg, 0.33 mmol) and 1-Boc-amino-3,6,9-trioxaundecanyl-11-bromide (47 mg, 0.13 mmol). The resulting mixture was stirred at room temperature for 72 h. The reaction mixture was diluted with water (1 mL) and extracted with EtOAc (2 mL x 3). The organic layers were then combined, washed with brine, dried over $Na_2SO_4$ and filtered. The resulting solution was concentrated *in vacuo* to afford the crude product which was further purified by flash column chromatography on silica gel (0-5% MeOH/$CH_2Cl_2$) to give **19** as a brown gum (25 mg, 29%). $^1$H NMR (400 MHz, $CDCl_3$) δ 7.55 – 7.36 (m, 1H), 7.18 (d, *J* = 8.1 Hz, 2H), 7.12 – 6.98 (m, 1H), 6.95 – 6.73 (m, 4H), 5.29 (s, residual DCM), 5.12 – 5.07 (m, 1H), 4.41 (s, 2H), 4.21 – 4.13 (m, 2H), 4.11 (q, J = 7.2, residual EtOAc), 4.03 – 3.94 (m, 2H), 3.90 – 3.85 (m, 2H), 3.80 – 3.59 (m, 12H), 3.57 – 3.48 (m, 2H), 3.35 – 3.26 (m, 2H), 3.04 – 2.96 (m, 1H), 2.95 (s, residual DMF), 2.88 (s, residual DMF), 2.04 (s, residual EtOAc), 2.01 – 1.84 (m, 4H), 1.75 – 1.61 (m, 4H), 1.47 – 1.41 (m, 18H). 1.24 (t, J = 7.1, residual EtOAc).

Step 6: 4N HCl in dioxane (4M, 100 µL) was added to a stirring solution of **19** (25 mg, 0.032 mmol) in MeOH (100 µL). The resulting solution was stirred at room temperature for 2 h, at which point the reaction mixture was concentrated *in vacuo* to afford **8** (20 mg, quantitative) as a brown gum. $^1$H NMR (600 MHz, DMSO) δ 9.42 (s, 2H), 8.60 (d, *J* = 7.9 Hz, 1H), 8.16 (s, 3H), 7.68 (d, *J* = 9.0 Hz, 1H), 7.47 – 7.42 (m, 2H), 7.31 – 7.19 (m, 1H), 7.24 (d, *J* = 2.4 Hz, 1H), 7.14 (dd, *J* = 9.0, 2.4 Hz, 1H), 7.08 (d, *J* = 8.8 Hz, 1H), 6.97 – 6.92 (m, 2H), 4.22 – 4.15 (m, 2H), 4.09 – 4.00 (m, 3H), 3.81 – 3.77 (m, 2H), 3.76 – 3.73 (m, 3H), 3.73 – 3.68 (m, 2H), 3.64 – 3.27 (m, 11H), 2.96 – 2.88 (m, 2H), 2.32 – 2.22 (m, 2H), 2.02 – 1.94 (m, 2H), 1.73 – 1.61 (m, 4H). $^{13}$C NMR (151 MHz, DMSO) δ 164.26, 159.76, 156.89, 156.06, 131.88, 131.71, 125.00, 123.31, 115.66, 114.62, 114.01, 96.99, 69.97, 69.81, 69.68, 69.67, 68.85, 68.10, 66.60, 55.25, 49.44, 46.63, 43.91, 38.46, 33.97, 28.47, 24.60. HRMS (ESI): calculated for ($C_{31}H_{45}N_5O_6$) $[M+H]^+$: 584.3443, found: 584.3434.

**Compound 9: PFI-7**; *N*-((1s,4s)-4-(1H-benzo[d]imidazol-2-yl)cyclohexyl)-2-(((1H-indol-2-yl)methyl)amino)acetamide

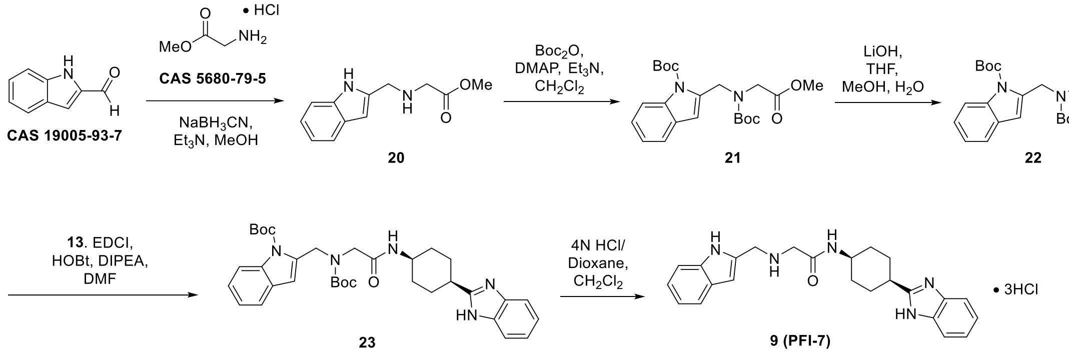


Step 1: To a solution of glycine methyl ester hydrochloride (16.0 g, 127 mmol), 1H-indole-2-carbaldehyde (7.40 g, 50.98 mmol) and triethylamine (7.57 mL, 56. 1 mmol) in methanol (60 mL) was added sodium cyanoborohyride (3.52 g, 56.1 mmol) at 0 °C. The resulting mixture was left to warm up to room temperature and stirred for 16 hours. The reaction mixture was diluted with

distilled water (60 mL) and extracted with EtOAc (60 mL x 4). The combined organic layers were washed with brine and dried over $Na_2SO_4$, filtered, and concentrated *in vacuo*. CombiFlash Silica gel column chromatography (0-0.5% MeOH/$CH_2Cl_2$) afforded **20** as a brown solid (5 g, 44.9%) and as a brown oil (3 g, 27%). LCMS and 1H NMR ($CDCl_3$) analysis confirmed the two batches contained the desired product and was combined for the next step. LCMS m/z 219 $[M+H]^+$. 1H NMR (400 MHz, CDCl3) δ 8.78 (s, 1H), 7.58 (dd, *J* = 7.9, 1.2 Hz, 1H), 7.35 (dd, *J* = 8.0, 1.1 Hz, 1H), 7.18 (t, *J* = 8.1, 7.1, 1.3 Hz, 1H), 7.11 (td, *J* = 7.4, 7.0, 1.1 Hz, 1H), 6.38 (dd, *J* = 2.0, 1.0 Hz, 1H), 4.02 (s, 2H), 3.76 (s, 3H), 3.46 (s, 2H).

Step 2: To a solution of **20** (3 g, 13.7 mmol) and $Boc_2O$ (10.5 g, 48.1 mmol) in $CH_2Cl_2$ (60 mL) was added DMAP (336 mg, 2.75 mmol) and triethylamine (4.17 g, 41.2 mmol). The resulting mixture was stirred at room temperature for 16 hours, whereupon another equivalent of $Boc_2O$ (3 g, 13.7 mmol) and DMAP (168 mg, 1.37 mmol) were added. After additional 16 h of stirring, the reaction mixture was concentrated *in vacuo* and diluted with aq. $NH_4Cl$ (100 mL). The aqueous solution was extracted with EtOAc (80 mL x 4), and the combined organic layers were dried over Na2SO4, filtered, and concentrated *in vacuo*. CombiFlash silica gel column chromatography (0-12.6% EtOAc/petroleum ether) afforded **21** as a yellow gum (4 g, 69.5%). LCMS m/z 441 $[M+Na]^+$. 1H NMR (400 MHz, CDCl3) δ 8.09 (dd, *J* = 8.3, 5.2 Hz, 1H), 7.50 (t, *J* = 6.5 Hz, 1H), 7.32 – 7.19 (m, 2H), 6.48 (d, *J* = 18.2 Hz, 1H), 4.91 (d, *J* = 14.2 Hz, 2H), 4.12 (s, 1H), 4.02 (s, 1H), 3.76 (d, *J* = 2.9 Hz, 3H), 1.71 (d, *J* = 7.2 Hz, 9H), 1.48 (d, *J* = 11.7 Hz, 9H).

Step 3: To a solution of **21** (4 g, 9.56 mmol) in MeOH (30 mL), THF (30 mL), and water (10 mL) at 0 °C was added lithium hydroxide (401 mg, 9.56 mmol) before removing the ice bath. After stirring at room temperature for 3 hours, the reaction mixture was concentrated *in vacuo* to afford **22** as a yellow gum (11.7 g, quantitative). This material was used directly, with no further purification, in the following step. LCMS m/z 427 $[M+Na]^+$ on crude reaction product.

Step 4: To a solution of **13** (6.23 g, 28.9 mmol) and **22** (11.7 g, 28.9 mmol) in DMF (80 mL) at 0° C was added HOBt (5.86 g, 43.4 mmol), EDCI (8.31 g, 43.4 mmol), and Hunig's base (14.2 mL, 86.7 mmol). The resulting mixture was warmed to room temperature and left to stir for 16 hours. The reaction mixture was diluted with aq. $NH_4Cl$ (100 mL) and extracted with EtOAc (80 mL x 4). The combined organic layers were washed with brine, dried over $Na_2SO_4$, filtered, and concentrated *in vacuo*. CombiFlash silica gel column chromatography (0-0.6% MeOH/$CH_2Cl_2$) afforded **23** as a light pink gum (12.0 g, 69%). LCMS m/z 602 $[M+H]^+$. 1H NMR (400 MHz, CDCl3) δ 8.08 – 7.95 (m, 2H w/CH from DMF), 7.45 – 7.39 (m, 3H), 7.25 – 7.11 (m, 4H), 6.37 (s, 1H), 4.87 (s, 2H), 4.10 – 3.97 (m, 1H), 3.93 (s, 2H), 3.08 – 2.96 (m, 1H), 2.12 – 1.93 (m, 2H), 1.93 – 1.82 (m, 2H), 1.81 – 1.66 (m, 1H), 1.63 (s, 9H), 1.39 (s, 12H).

Step 5: To a solution of **23** (200 mg, 0.332 mmol) in $CH_2Cl_2$ (5 mL) was added a solution of HCl in 1,4-dioxane (4 M, 1 mL). The resulting mixture was stirred at room temperature for 3 hours before adding additional solution of HCl in 1,4-dioxane (4 M, 0.5 mL). The reaction mixture was then brought to 40 °C and stirred for 5.5 hours before concentrating *in vacuo* to afford **9** as a pink solid (160 mg, quantitative). $^1$H NMR (600 MHz, DMSO) δ 15.39 (s, 2H), 11.36 (s, 1H), 9.67 (s, 2H), 8.56 (d, *J* = 7.8 Hz, 1H), 7.82 – 7.76 (m, 2H), 7.55 – 7.50 (m, 3H), 7.39 (d, *J* = 8.2 Hz, 1H), 7.10 (t, *J* = 7.6 Hz, 1H), 6.99 (t, *J* = 7.4 Hz, 1H), 6.61 (s, 1H), 4.37 (s, 2H), 4.05 – 4.01 (m, 1H), 3.77 (s, 2H), 3.38 – 3.31 (m, 1H), 2.32 – 2.22 (m, 2H), 2.06 – 1.95 (m, 2H), 1.75 – 1.60 (m, 4H). $^{13}$C NMR (151 MHz, DMSO) δ 164.17, 156.68, 136.22, 130.67,

128.91, 127.44, 125.56, 121.98, 120.31, 119.36, 113.73, 111.51, 103.83, 46.62, 43.97, 43.22, 33.95, 28.39, 24.53. HRMS (ESI): calculated for ($C_{24}H_{27}N_5O$) $[M+H]^+$: 402.2288, found: 402.2283.

**Compound 10**: **PFI-7N**; N-((1s,4s)-4-(1H-benzo[d]imidazol-2-yl)cyclohexyl)-2-(N-((1H-indol-2-yl)methyl)acetamido)acetamide

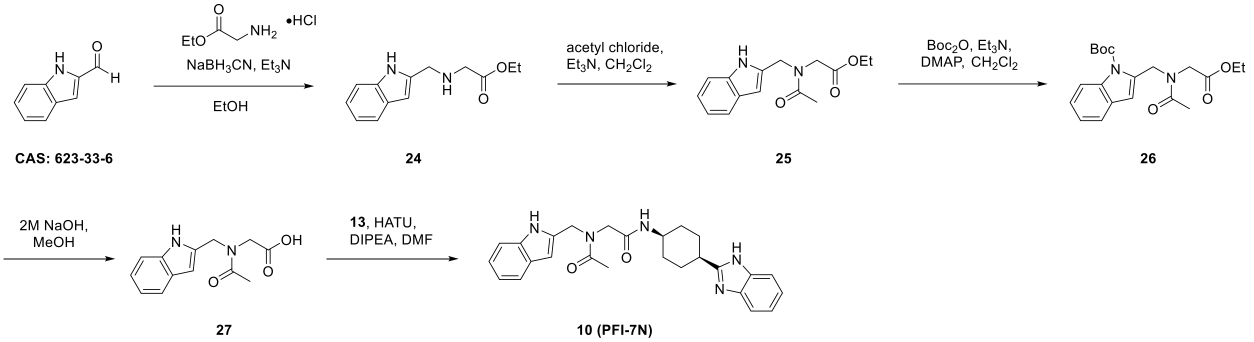


Step 1: To a solution of glycine ethyl ester hydrochloride (1.44 g, 10.3 mmol), 1H-indole-2-carbaldehyde (1.0 g, 6.89 mmol) and triethylamine (1.44 mL, 10.3 mmol) in ethanol (12 mL) was added sodium cyanoborohyride (649 mg, 10.3 mmol) at 0 °C. The resulting mixture was left to warm up to room temperature and stirred for 16 hours. The reaction mixture was diluted with distilled water (10 mL) and extracted with EtOAc (15 mL x 3). The combined organic layers were washed with $NaHCO_3$, brine, and dried over $Na_2SO_4$, filtered, and concentrated *in vacuo*. CombiFlash Silica gel column chromatography (0 → 50% EtOAc/petroleum ether) afforded **24** as a brown solid (970 mg, 61%). $^1$H NMR (400 MHz, $CDCl_3$) δ 8.65 (s, 1H), 7.59 – 7.52 (m, 1H), 7.37 – 7.30 (m, 1H), 7.15 (ddd, *J* = 8.2, 7.1, 1.3 Hz, 1H), 7.08 (ddd, *J* = 8.1, 7.1, 1.1 Hz, 1H), 6.38 – 6.33 (m, 1H), 5.30 (s, residual DCM), 4.20 (q, *J* = 7.2 Hz, 2H), 4.00 (s, 2H), 3.42 (s, 2H), 2.06 (s, 1H), 1.27 (t, *J* = 7.1 Hz, 3H).

Step 2: To a solution of **24** (200 mg, 0.86 mmol) in anhydrous $CH_2Cl_2$ (4.5 mL) was added triethylamine (0.18 mL, 1.29 mmol) followed by dropwise addition of acetyl chloride (67.6 mg, 0.86 mmol) in anhydrous $CH_2Cl_2$ (0.5 mL). The resulting mixture was left to stir at room temperature for 16h. The reaction mixture was diluted with water (10 mL) and extracted with EtOAc (15 mL x 3). The organic layers were combined, washed with brine, and dried over $Na_2SO_4$. It was then filtered and concentrated *in vacuo* to afford the crude product which was further purified *via* flash column chromatography on silica gel (0 → 70% EtOAc/petroleum ether) to give **25** as a yellow gum (236 mg, quantitative). $^1$H NMR (400 MHz, $CDCl_3$) δ 8.88 (s, 1H), 7.59 – 7.50 (m, 1H), 7.35 – 7.28 (m, 1H), 7.21 – 7.03 (m, 2H), 6.40 – 6.34 (m, 1H), 5.30 (s, residual DCM), 4.78 (s, 1H), 4.62 (s, 1H), 4.27 (q, *J* = 7.1 Hz, 1H), 4.17 (s, 1H), 4.12 (q, *J* = 7.2 Hz, 1H), 4.02 (s, 1H), 2.14 – 2.06 (m, 3H), 1.58 (s, residual $H_2O$), 1.31 (t, *J* = 7.2 Hz, 1H), 1.19 (t, *J* = 7.1 Hz, 2H).

Step 3: To a solution of **25** (236 mg, 0.86 mmol) in $CH_2Cl_2$ (5 mL) was added triethylamine (0.36 mL, 2.58 mmol) followed by $Boc_2O$ (375 mg, 1.72 mmol) and DMAP (21 mg, 0.17 mmol). The resulting solution was stirred at room temperature for 16 h. The reaction mixture was diluted with water (10 mL) and extracted with EtOAc (10 mL x 3) before combining the organic layers. The organic layer was then washed with brine, dried over $Na_2SO_4$, filtered, and concentrated *in vacuo*. Flash column chromatography on silica gel (0 → 25% EtOAc/petroleum ether) afforded **26** as a yellow gum (193 mg, 60%). $^1$H NMR (400 MHz, $CDCl_3$) δ 8.10 – 8.01 (m, 1H), 7.53 – 7.45 (m, 1H), 7.33 – 7.17 (m, 2H), 6.50 – 6.45 (m, 1H), 5.30 (s, residual DCM), 5.03 – 4.94 (m, 2H), 4.23 – 4.16 (m, 4H), 2.21 (s, 3H), 1.70 (s, 9H), 1.57 (s, residual $H_2O$), 1.29 – 1.23 (m, 3H).

Step 4: 2M NaOH (0.26 mL, 0.51 mmol) was added to a solution of **26** (193 mg, 0.51 mmol) in MeOH (2 mL). The resulting mixture was left to stir at room temperature for 16h upon which TLC analysis (5:1 petroleum ether:EtOAc) showed complete conversion of the starting material. The reaction mixture was diluted with water (10 mL) and adjusted to pH ~3 with 10% citric acid (aqueous). The solution was then diluted with EtOAc (15 mL) and the resulting layers were separated. The aqueous layer was extracted with EtOAc (15 mL x 2) before combining the organic layers. Washed the organic layer with brine, dried over $Na_2SO_4$, filtered, and concentrated in vacuo to afford **27** as a yellow solid (108 mg, 86%). $^1$H NMR (400 MHz, $CDCl_3$) δ 8.95 (s, 1H), 7.59 – 7.50 (m, 1H), 7.38 – 7.27 (m, 1H), 7.22 – 7.02 (m, 2H), 6.42 – 6.36

(m, 1H), 4.77 (s, 1H), 4.65 (s, 1H), 4.18 (s, 1H), 4.12 (q, J = 7.1, residual EtOAc), 4.06 (s, 1H), 2.16 – 2.09 (m, 3H), 2.05 (s, residual EtOAc), 1.26 (t, J = 7.2 , residual EtOAc).

Step 5: To a solution of **27** (108 mg, 0.44 mmol) in anhydrous DMF (3 mL) was added HATU (247 mg, 0.65 mmol) followed by **13** (152 mg, 0.53 mmol) and DIPEA (0.22 mL, 1.32 mmol). The resulting mixture was left to stir at room temperature for 16h. The crude reaction mixture was directly purified by reversed-phase HPLC (Column: C18, 30 x 150 mm, 5 µm; Mobile phase A: 10 mM aqueous ammonium bicarbonate solution containing 0.05% ammonium hydroxide; Mobile phase B: acetonitrile; Gradient 24% to 64% B; Flow rate: 30 mL/min) and dried by lyophilization to afford **10** as a white solid (90 mg, 46%). $^{1}$H NMR (500 MHz, DMSO) δ 12.10 (s, 1H), 11.29 (s, 1H), 10.85 (s, 1H), 7.94 (d, *J* = 7.7 Hz, 1H), 7.79 (d, *J* = 7.6 Hz, 1H), 7.56 – 7.48 (m, 1H), 7.46 – 7.38 (m, 2H), 7.31 (t, *J* = 8.3 Hz, 1H), 7.15 – 7.06 (m, 2H), 7.06 – 6.97 (m, 1H), 6.97 – 6.89 (m, 1H), 6.30 – 6.23 (m, 1H), 4.68 (s, 1H), 4.57 (s, 1H), 3.90 (d, *J* = 8.0 Hz, 2H), 3.87 – 3.80 (m, 1H), 3.34 (s, residual $H_2O$), 3.05 – 2.96 (m, 1H), 2.19 – 2.07 (m, 3H), 1.99 (s, 1H), 1.88 – 1.77 (m, 2H), 1.70 – 1.52 (m, 4H). $^{13}$C NMR (126 MHz, DMSO) δ 170.79, 170.33, 167.88, 167.30, 157.68, 143.20, 143.18, 136.37, 136.33, 135.45, 135.39, 134.34, 134.31, 127.87, 127.83, 121.42, 121.40, 120.96, 120.71, 119.72, 119.49, 118.97, 118.81, 118.19, 111.18, 111.12, 110.73, 100.31, 99.83, 50.06, 48.05, 46.90, 45.63, 42.84, 34.45, 28.84, 26.47, 26.43, 21.47, 21.43. HRMS (ESI): calculated for ($C_{26}H_{29}N_5O_2$) $[M+H]^+$: 444.2394, found: 444.2387.

### Library chemistry protocol for Scheme 1

A solution of N-[*cis*-4-(1H-benzimidazol-2-yl)cyclohexyl]-2-chloroacetamide (15 mmol, 1 equiv.) in DMSO (0.075 M) was treated with amine monomer (20 mmol, 1.3 equiv.) followed by Hunig's base (45 mmol, 3.0 equiv.). The solution was stirred in a vial at 70 $^{o}$C for 6 h or until determined to be complete by LCMS. The products were purified by preparative HPLC using reverse-phase HPLC on an Xbridge BEH column (10x100 mm, 5 um) eluting with either acetonitrile/water (0.225% formic acid) or acetonitrile/NH4OH (pH = 10). Flow rate = 4 mL/min.

**NMR Spectra of Final Compounds and Novel Intermediates**

**Compound 5:**

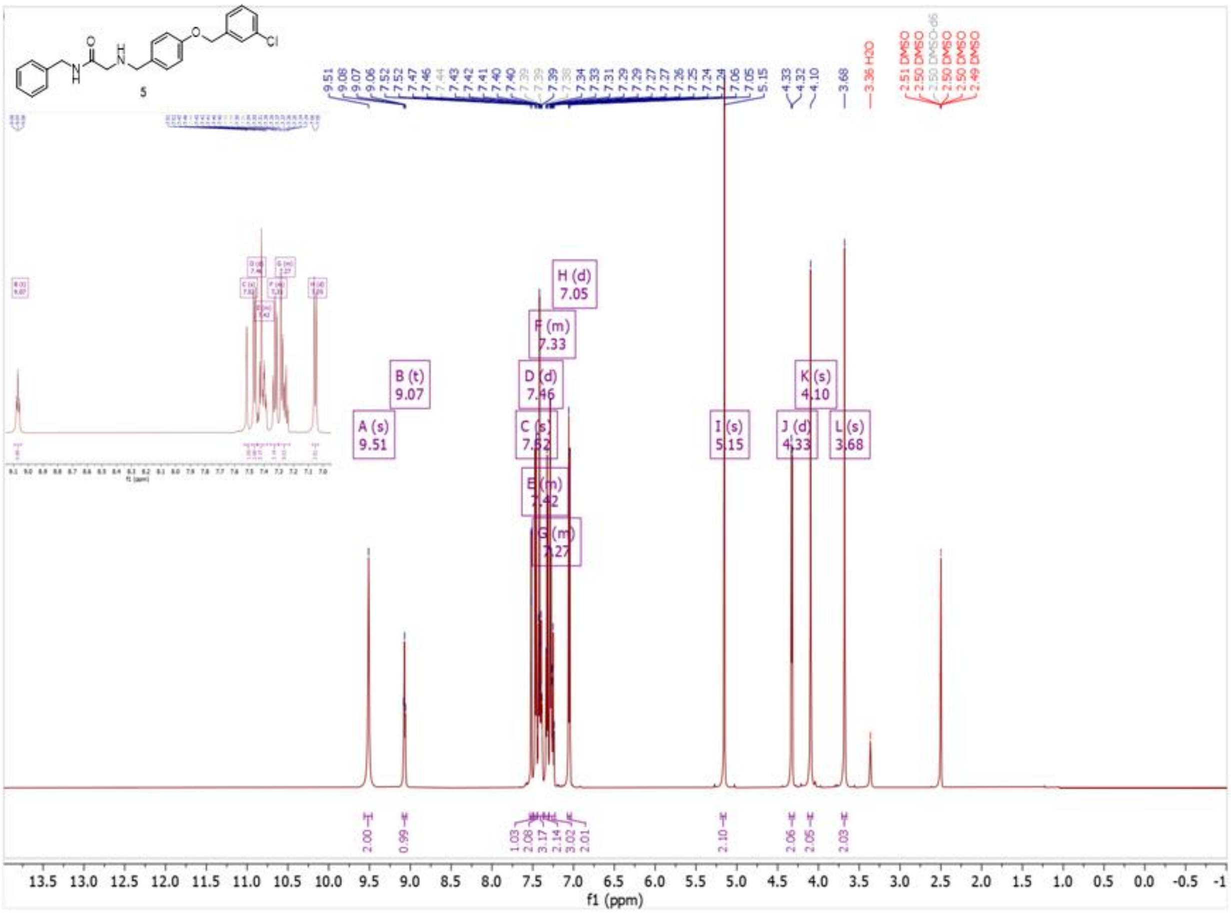

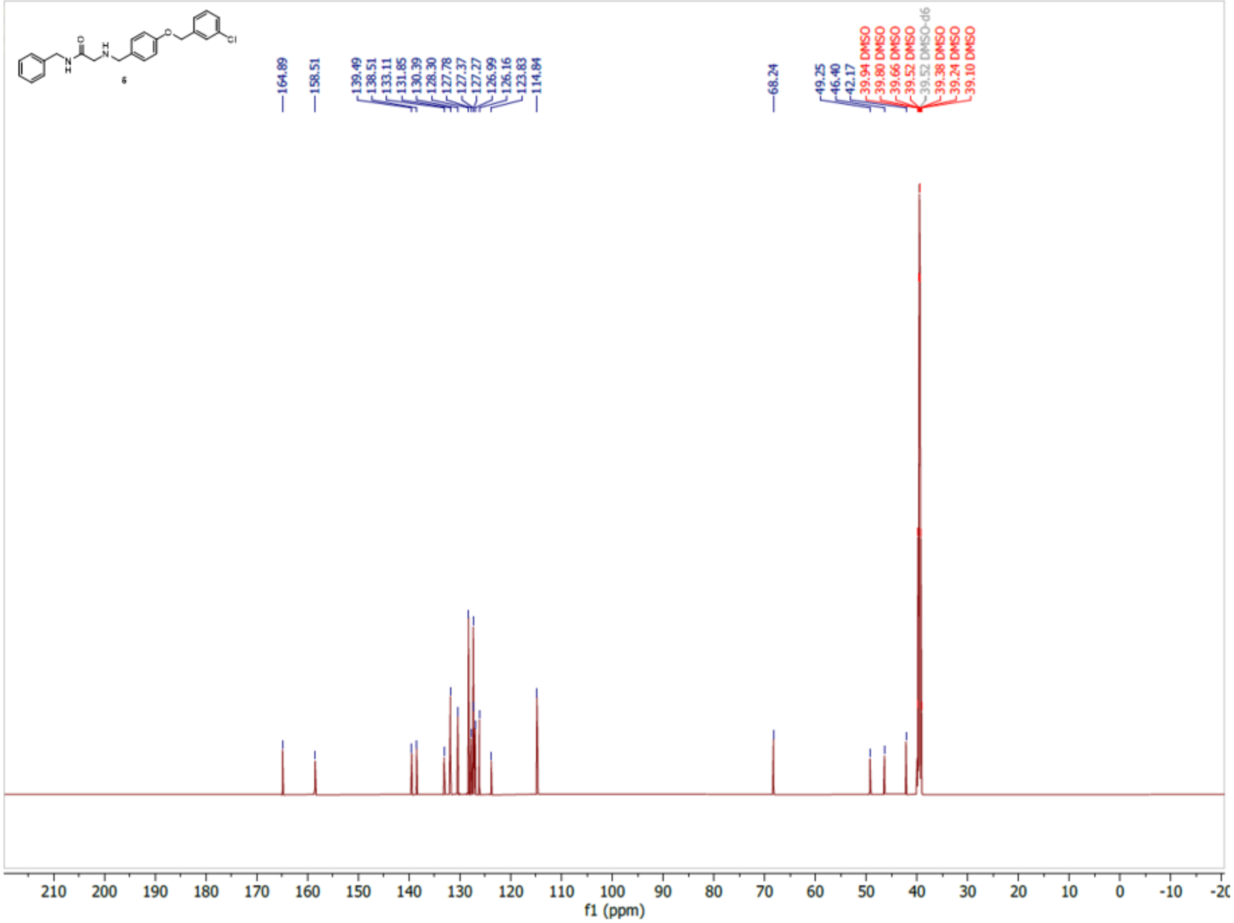
6
164.89
158.51
139.49
138.51
133.11
131.85
130.39
128.30
127.78
127.37
127.27
126.99
126.16
123.83
114.84
68.24
49.25
46.40
42.17
39.94 DMSO
39.80 DMSO
39.66 DMSO
39.52 DMSO
39.52 DMSO-d6
39.38 DMSO
39.24 DMSO
39.10 DMSO
f1 (ppm)

**Compound 6:**

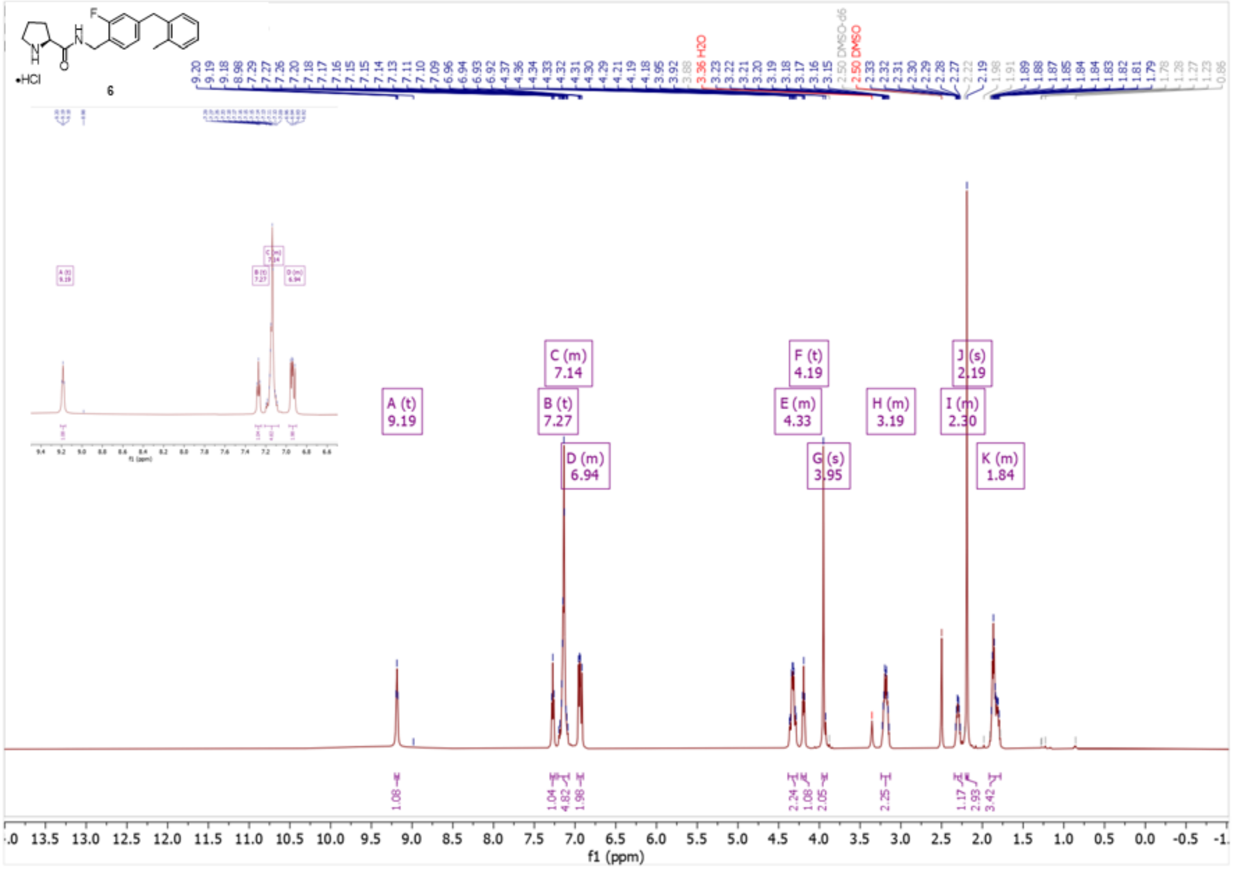
•HCl
6
A (t)
9.19
B (t)
7.27
C (m)
7.14
D (m)
6.94
E (m)
4.33
F (t)
4.19
G (s)
3.95
H (m)
3.19
I (m)
2.30
J (s)
2.19
K (m)
1.84
f1 (ppm)

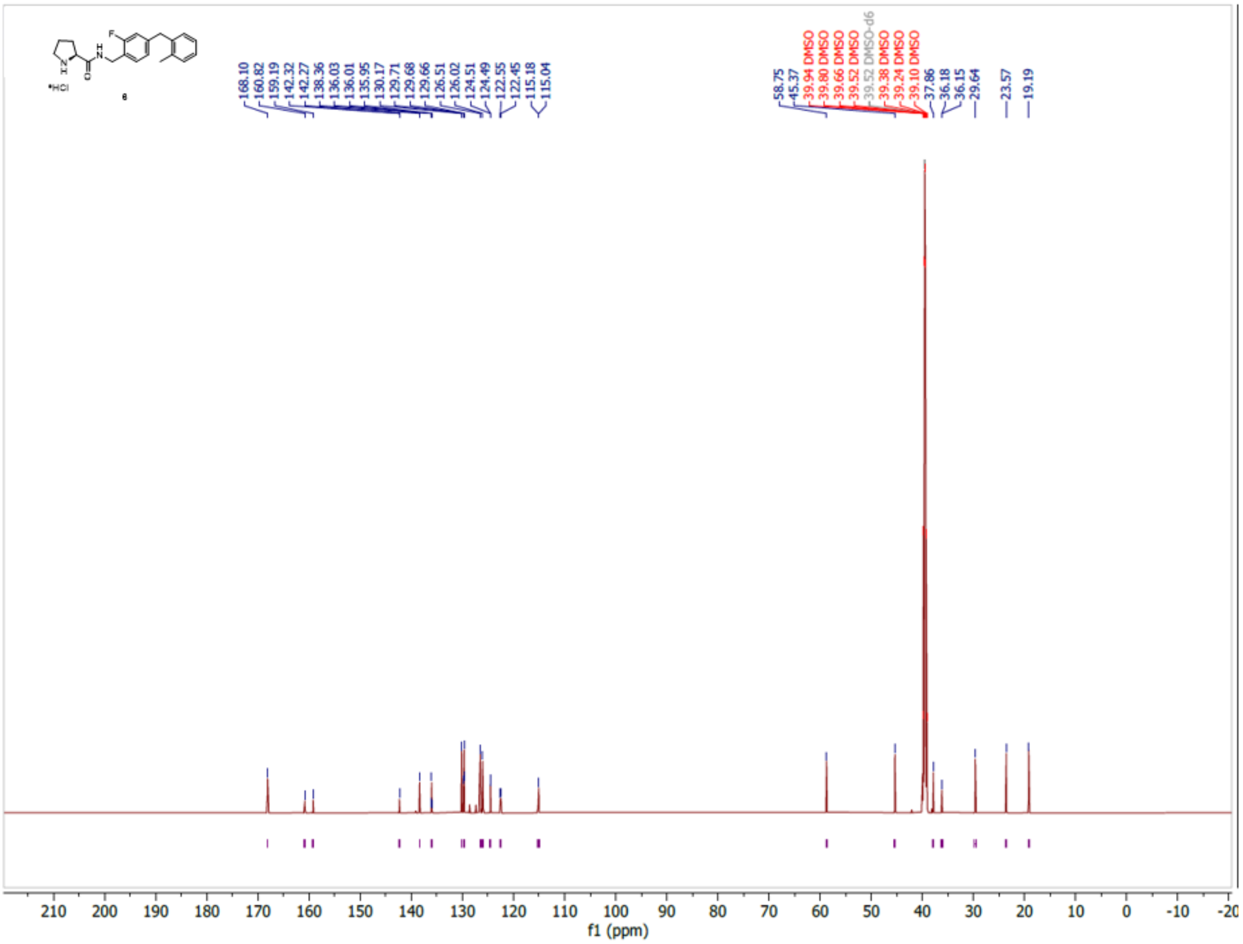

168.10
160.82
159.19
142.32
142.27
138.36
136.03
136.01
135.95
130.17
129.71
129.68
129.66
126.51
126.02
124.51
124.49
122.55
122.45
115.18
115.04
58.75
45.37
39.94 DMSO
39.80 DMSO
39.66 DMSO
39.52 DMSO
39.52 DMSO-d6
39.38 DMSO
39.24 DMSO
39.10 DMSO
37.86
36.18
36.15
29.64
23.57
19.19
210 200 190 180 170 160 150 140 130 120 110 100 90 80 70 60 50 40 30 20 10 0 -10 -20
f1 (ppm)

**Compound 7, PFI-E3H1:**

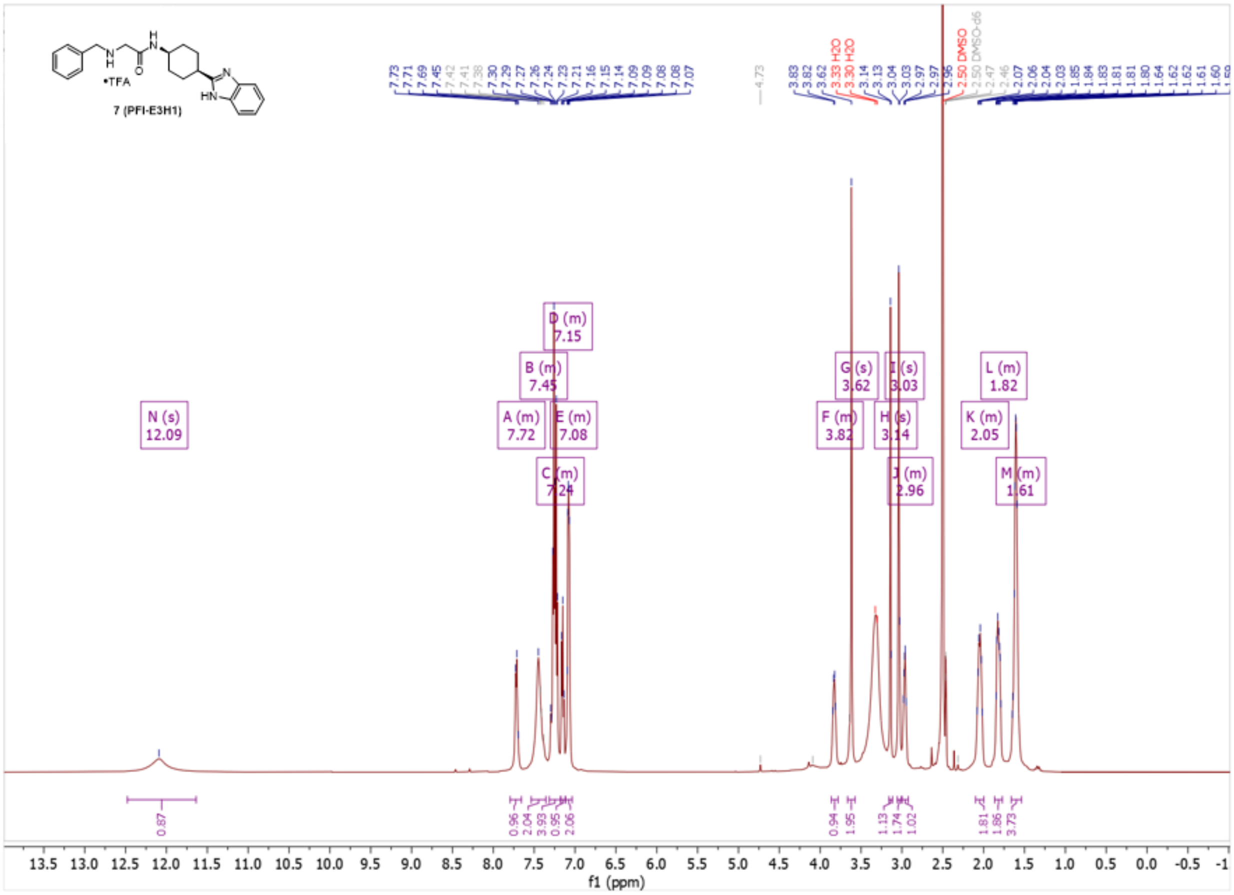

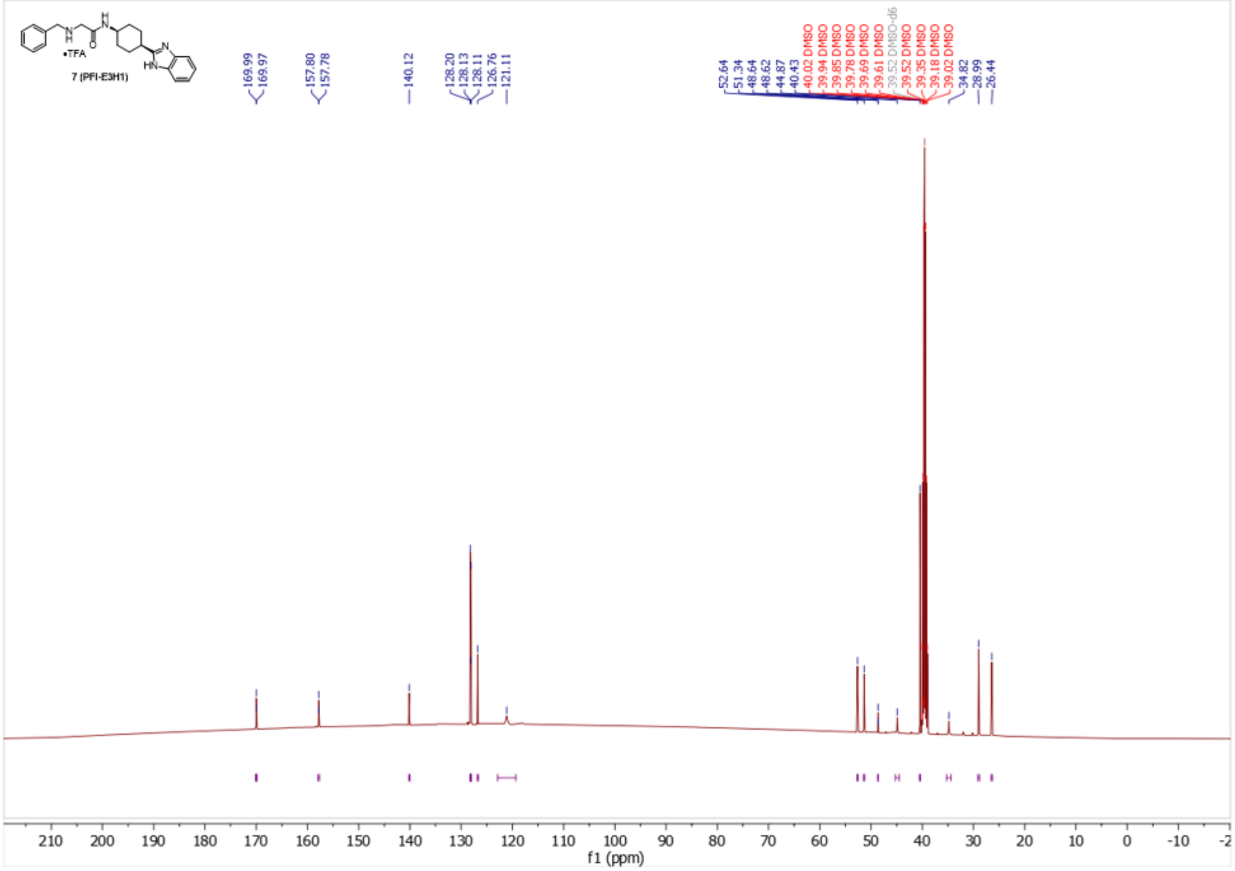
•TFA
7 (PFI-E3H1)
169.99
169.97
157.80
157.78
140.12
128.20
128.13
128.11
126.76
121.11
52.64
51.34
48.64
48.62
44.87
40.43
40.02 DMSO
39.94 DMSO
39.85 DMSO
39.78 DMSO
39.69 DMSO
39.61 DMSO
39.52 DMSO-d6
39.52 DMSO
39.35 DMSO
39.18 DMSO
39.02 DMSO
34.82
28.99
26.44
f1 (ppm)

**Compound 13:**

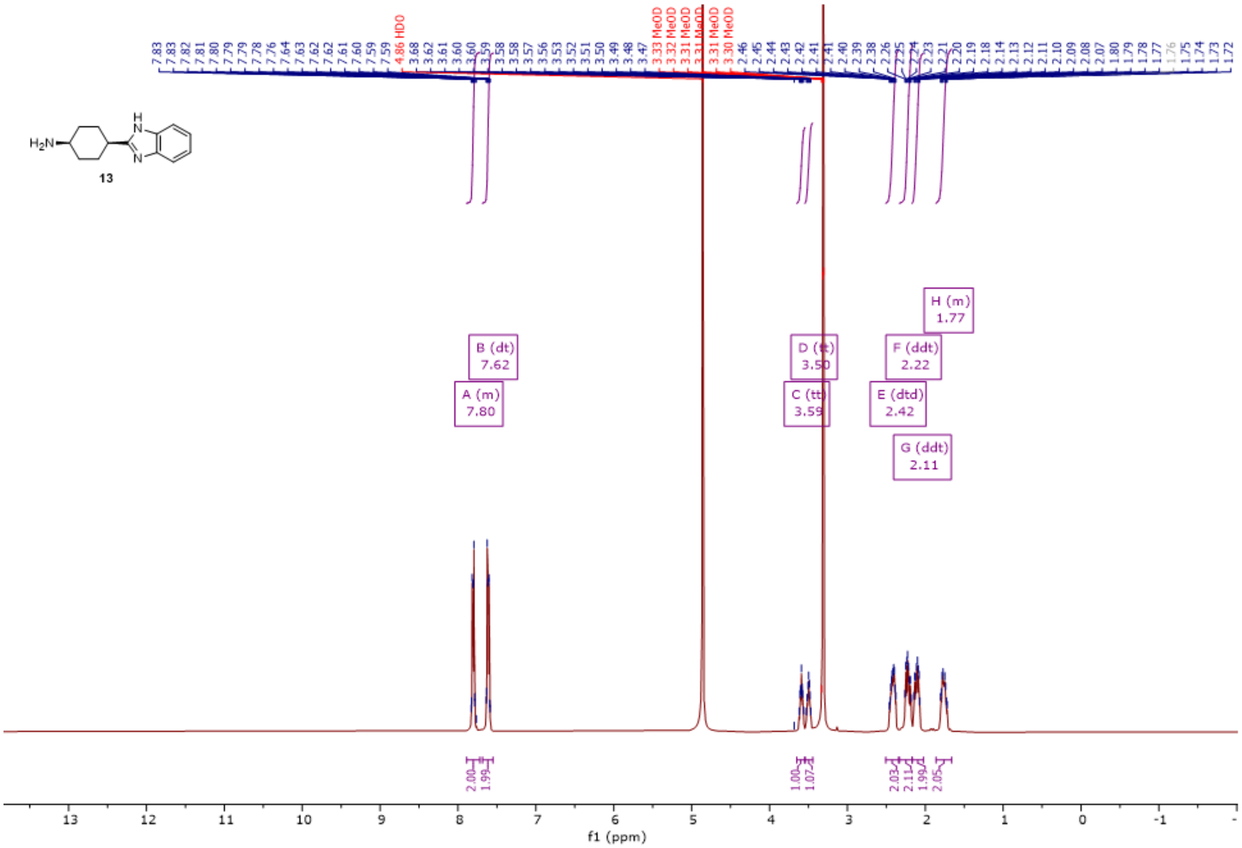

H2N
13
A (m)
7.80
B (dt)
7.62
C (tt)
3.59
D (tt)
3.50
E (dtd)
2.42
F (ddt)
2.22
G (ddt)
2.11
H (m)
1.77
4.86 HDO
f1 (ppm)

**Compound 15:**

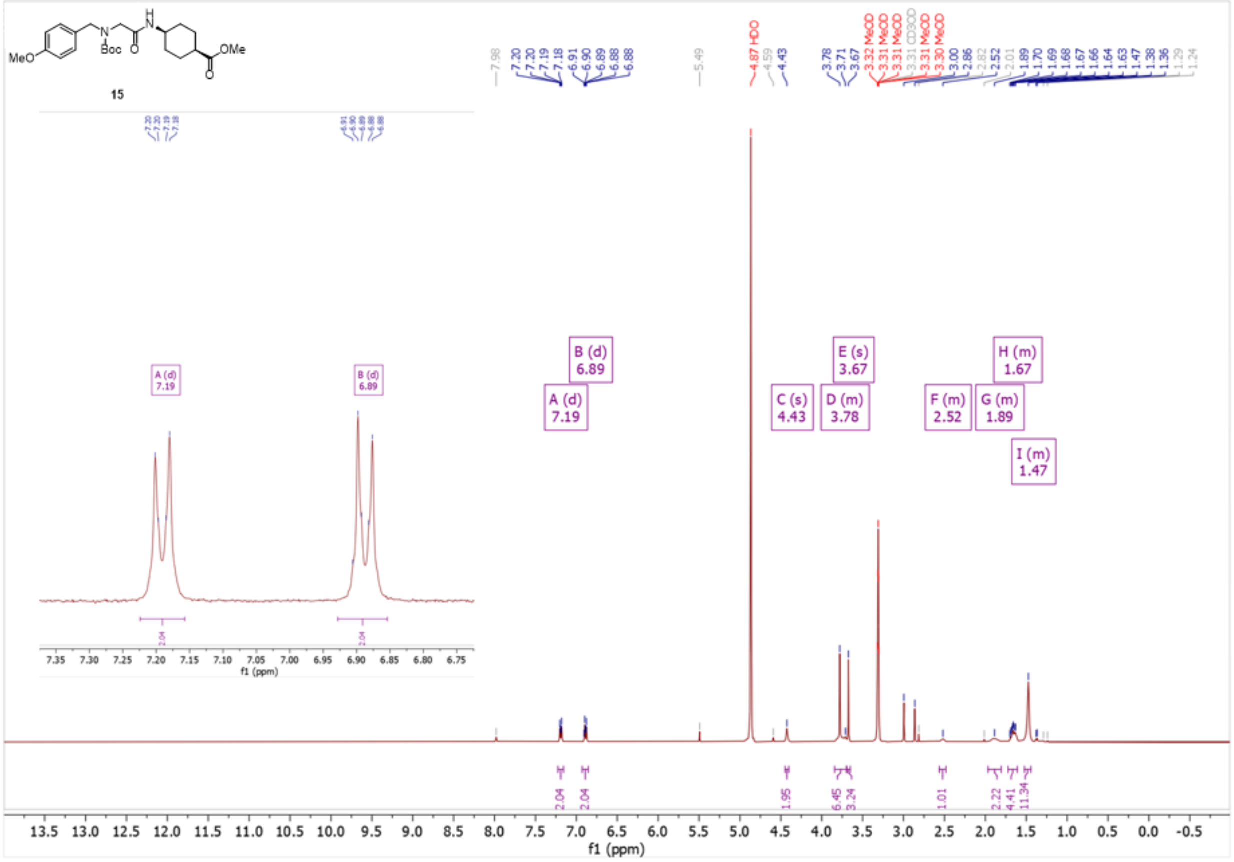

15
A (d)
7.19
B (d)
6.89
C (s)
4.43
D (m)
3.78
E (s)
3.67
F (m)
2.52
G (m)
1.89
H (m)
1.67
I (m)
1.47
f1 (ppm)

**Compound 16:**

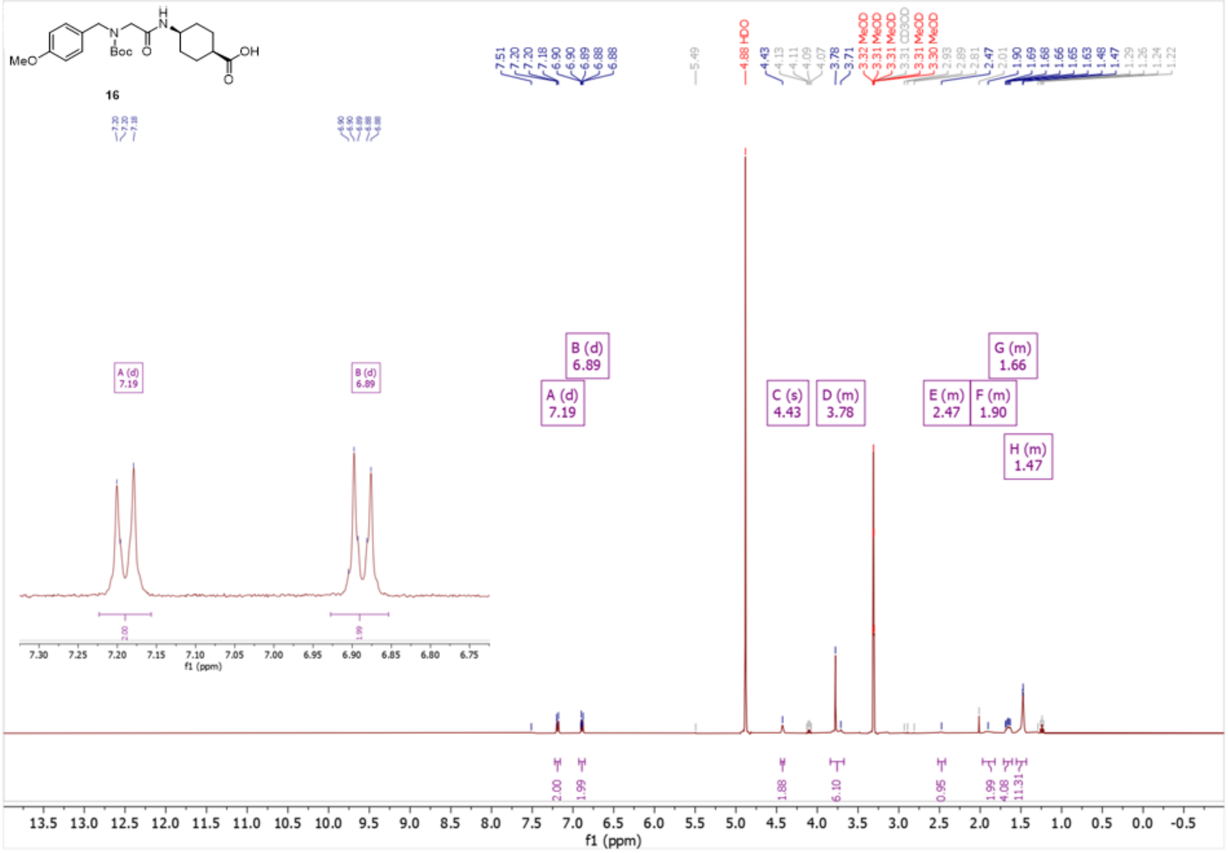

**Compound 17:**

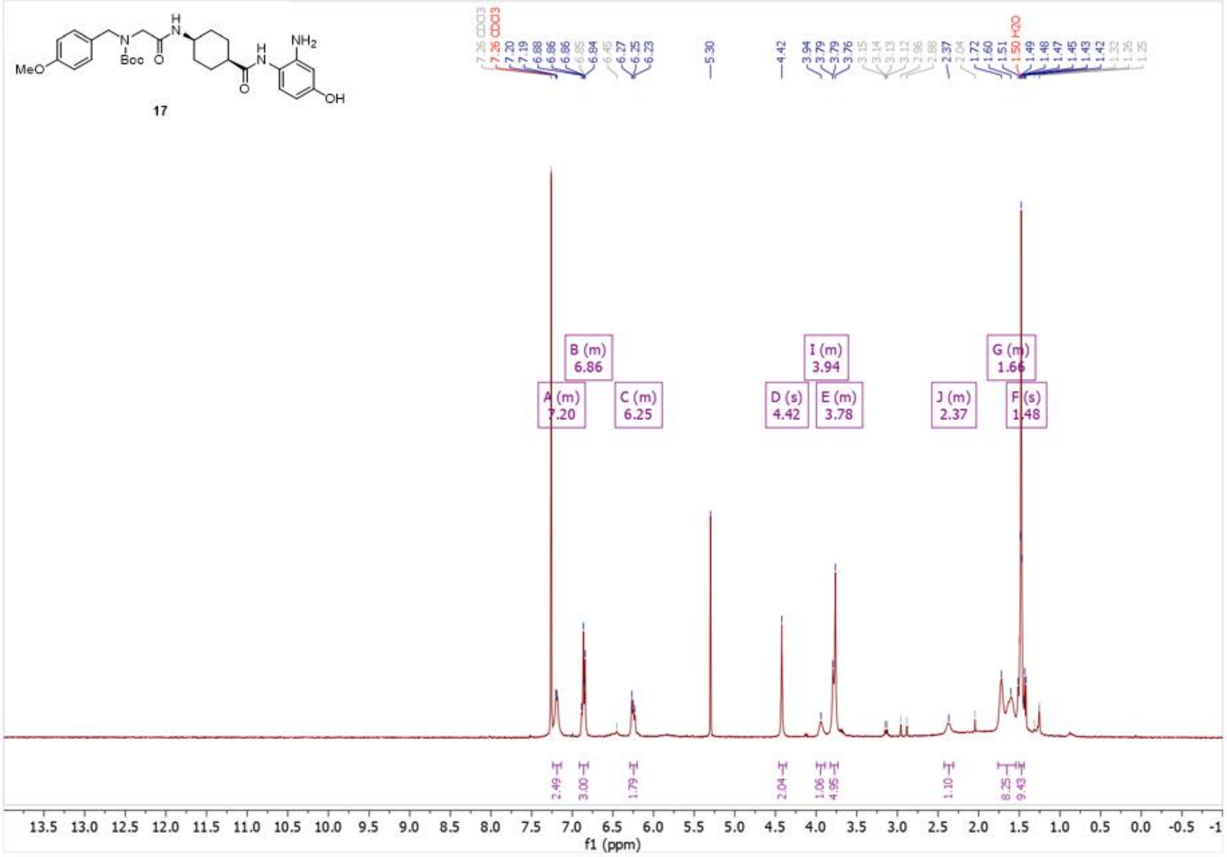

MeO
N
Boc
O
H
N
H
N
NH2
O
OH
17
7.26 CDCl3
7.26 CDCl3
7.20
7.19
6.88
6.86
6.86
6.85
6.84
6.45
6.27
6.25
6.23
5.30
4.42
3.94
3.79
3.79
3.76
3.15
3.14
3.13
3.12
2.96
2.88
2.37
2.04
1.72
1.60
1.51
1.50 H2O
1.49
1.48
1.47
1.45
1.43
1.42
1.32
1.26
1.25
B (m)
6.86
A (m)
7.20
C (m)
6.25
D (s)
4.42
I (m)
3.94
E (m)
3.78
J (m)
2.37
G (m)
1.66
F (s)
1.48
2.49
3.00
1.79
2.04
1.06
4.95
1.10
8.25
9.43
13.5 13.0 12.5 12.0 11.5 11.0 10.5 10.0 9.5 9.0 8.5 8.0 7.5 7.0 6.5 6.0 5.5 5.0 4.5 4.0 3.5 3.0 2.5 2.0 1.5 1.0 0.5 0.0 -0.5 -1
f1 (ppm)

**Compound 18:**

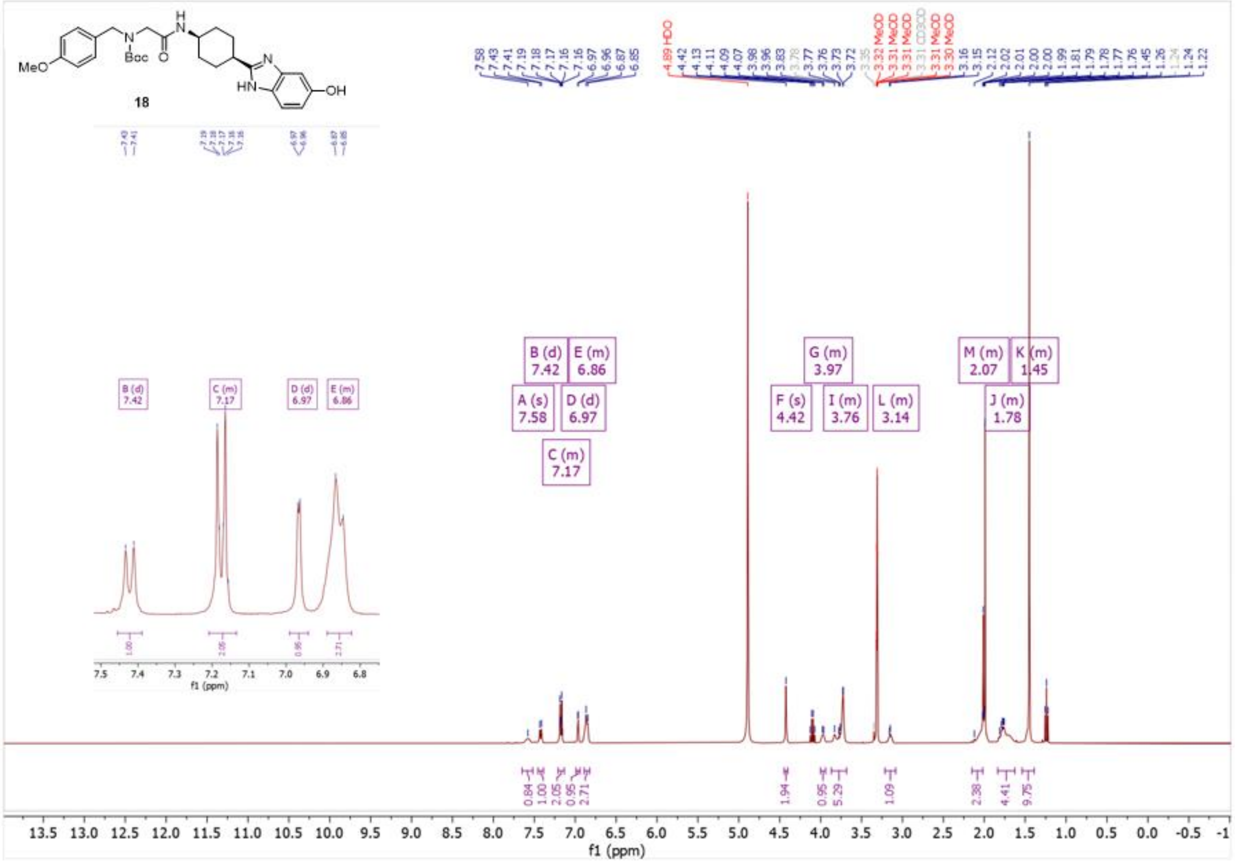

18
MeO
Boc
OH
HN
A (s) 7.58
B (d) 7.42
C (m) 7.17
D (d) 6.97
E (m) 6.86
F (s) 4.42
G (m) 3.97
I (m) 3.76
L (m) 3.14
M (m) 2.07
J (m) 1.78
K (m) 1.45
f1 (ppm)

**Compound 19:**

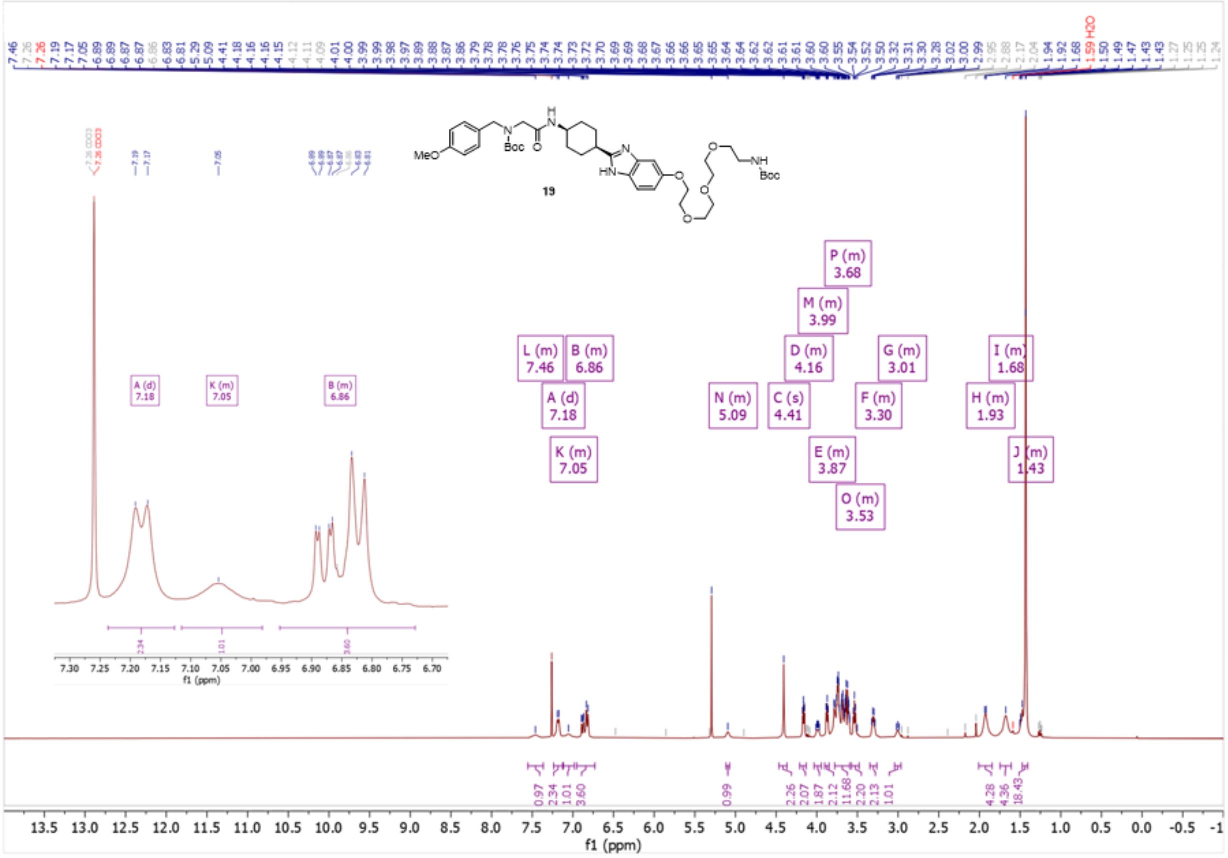

**Compound 8:**

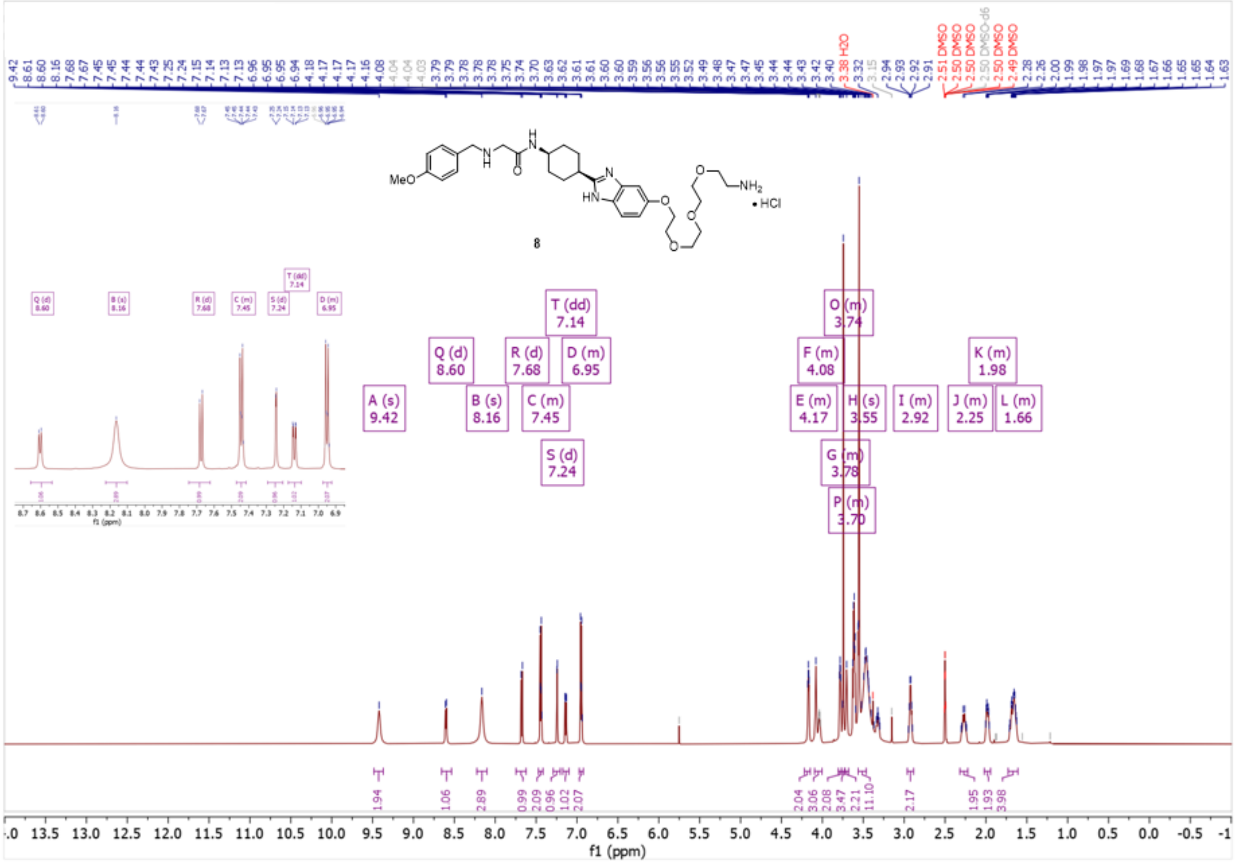

MeO
NH2
• HCl
8
A (s) 9.42
B (s) 8.16
C (m) 7.45
D (m) 6.95
E (m) 4.17
F (m) 4.08
G (m) 3.78
H (s) 3.55
I (m) 2.92
J (m) 2.25
K (m) 1.98
L (m) 1.66
O (m) 3.74
P (m) 3.70
Q (d) 8.60
R (d) 7.68
S (d) 7.24
T (dd) 7.14
3.38 H2O
2.50 DMSO
f1 (ppm)

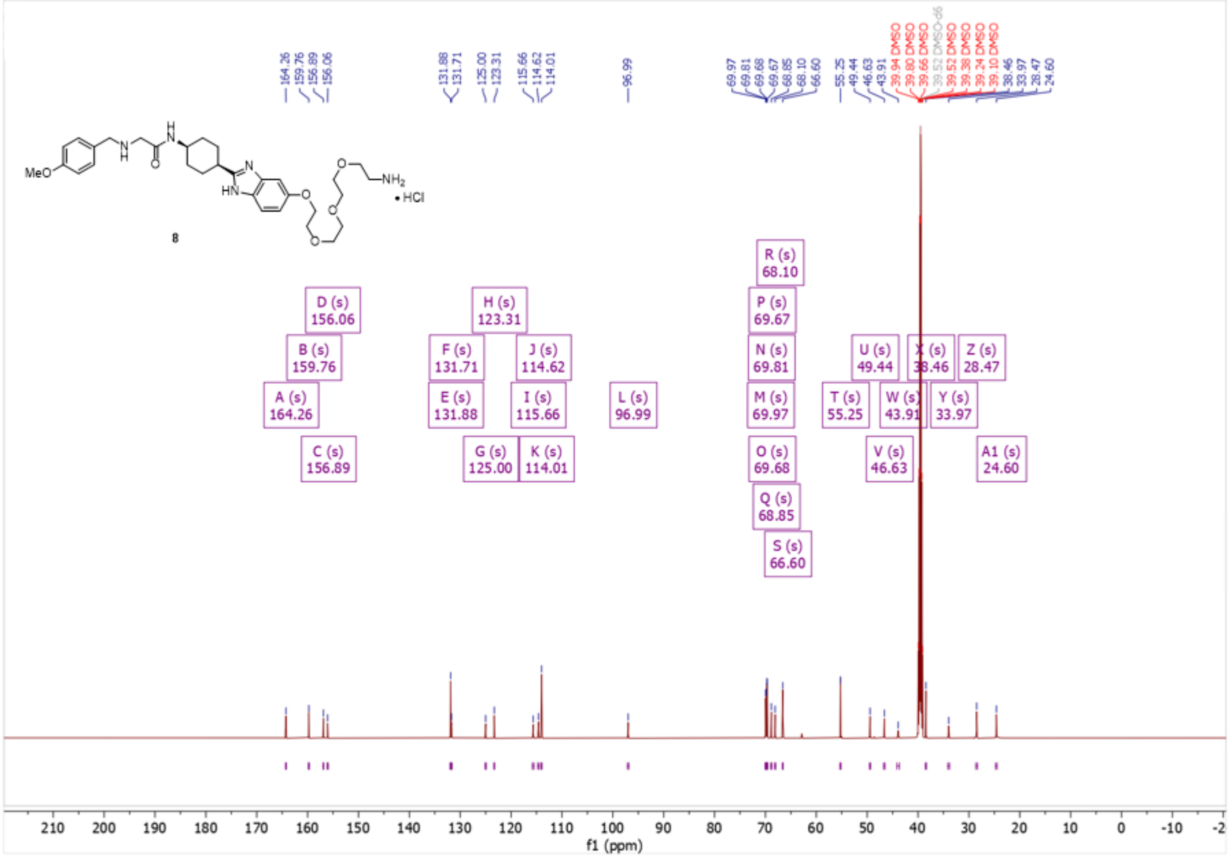
8
MeO
NH2
• HCl
f1 (ppm)
A (s) 164.26
B (s) 159.76
C (s) 156.89
D (s) 156.06
E (s) 131.88
F (s) 131.71
G (s) 125.00
H (s) 123.31
I (s) 115.66
J (s) 114.62
K (s) 114.01
L (s) 96.99
M (s) 69.97
N (s) 69.81
O (s) 69.68
P (s) 69.67
Q (s) 68.85
R (s) 68.10
S (s) 66.60
T (s) 55.25
U (s) 49.44
V (s) 46.63
W (s) 43.91
X (s) 38.46
Y (s) 33.97
Z (s) 28.47
A1 (s) 24.60

**Compound 20:**

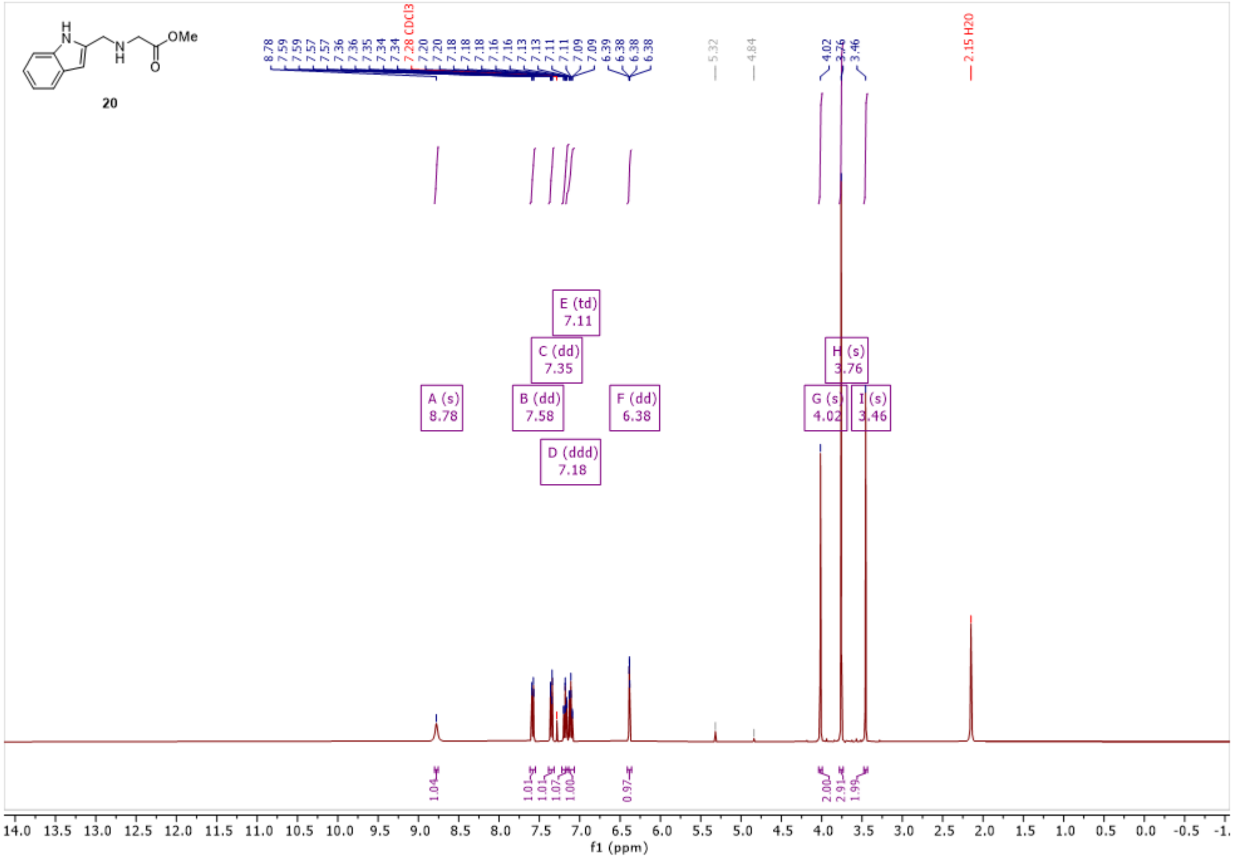

OMe
20
A (s) 8.78
B (dd) 7.58
C (dd) 7.35
D (ddd) 7.18
E (td) 7.11
F (dd) 6.38
G (s) 4.02
H (s) 3.76
I (s) 3.46
7.28 CDCl3
2.15 H2O
f1 (ppm)

**Compound 21:**

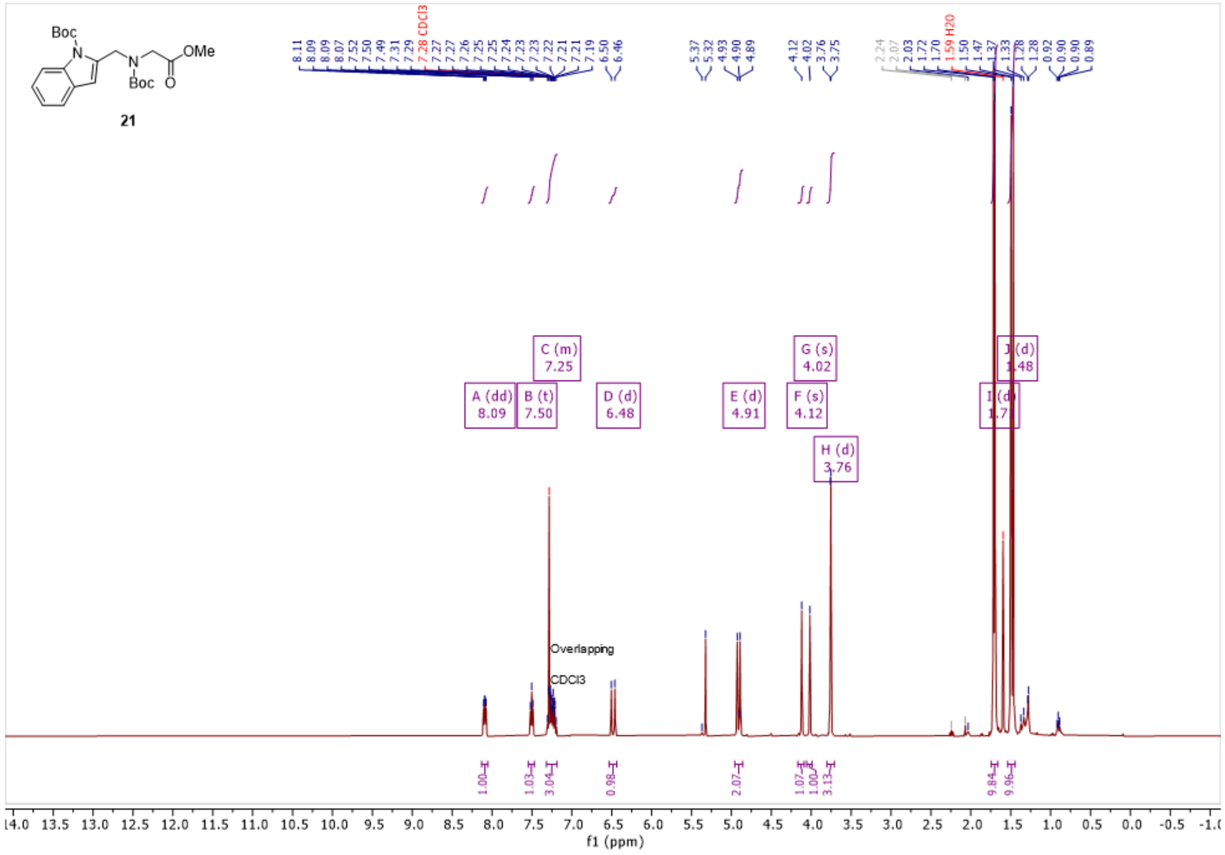

**Compound 23:**

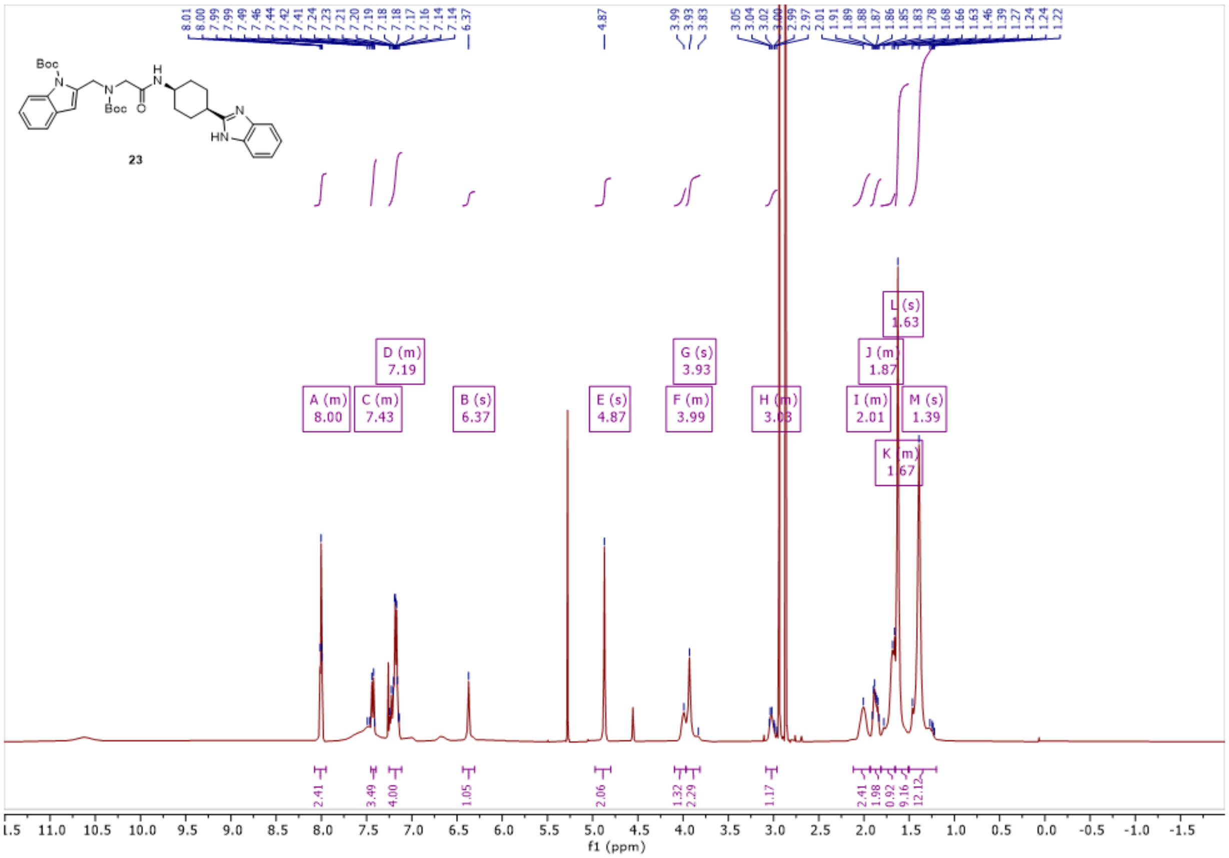

**Compound 9, PFI-7:**

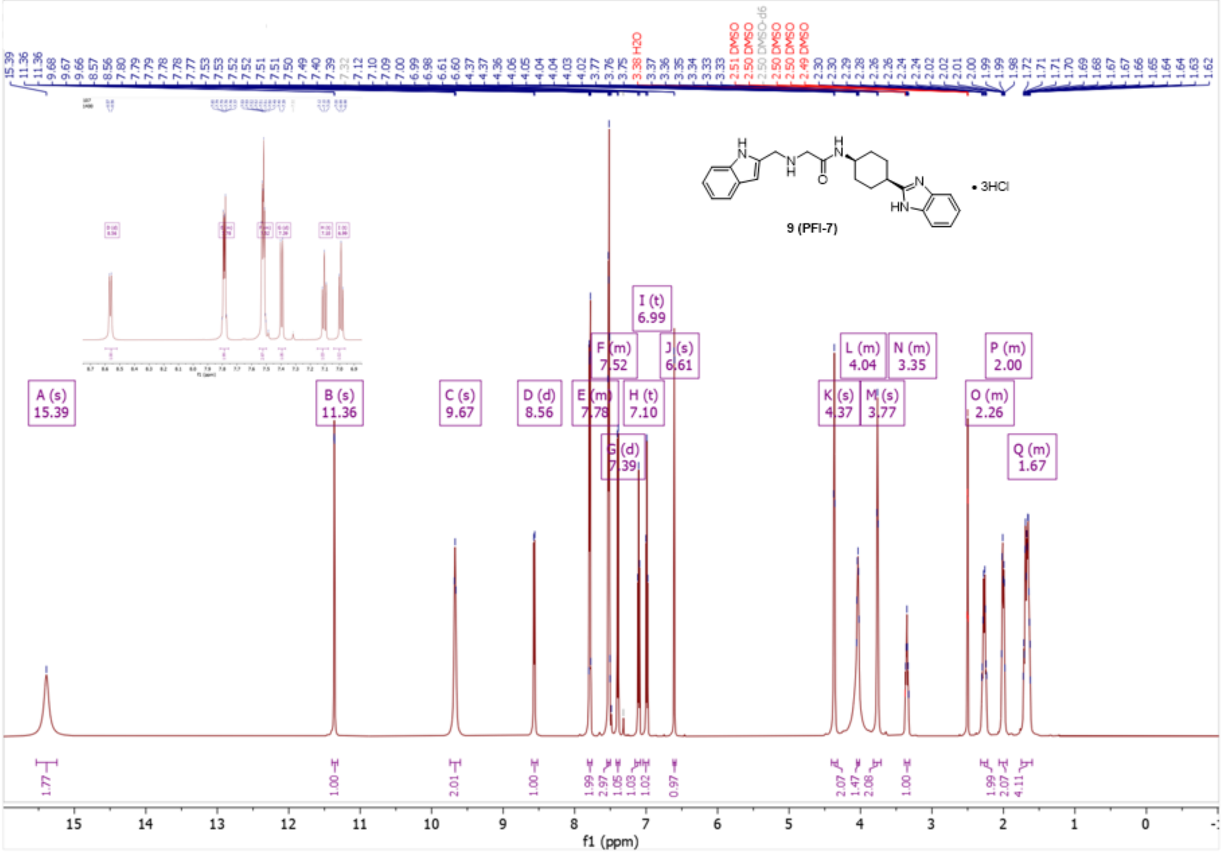

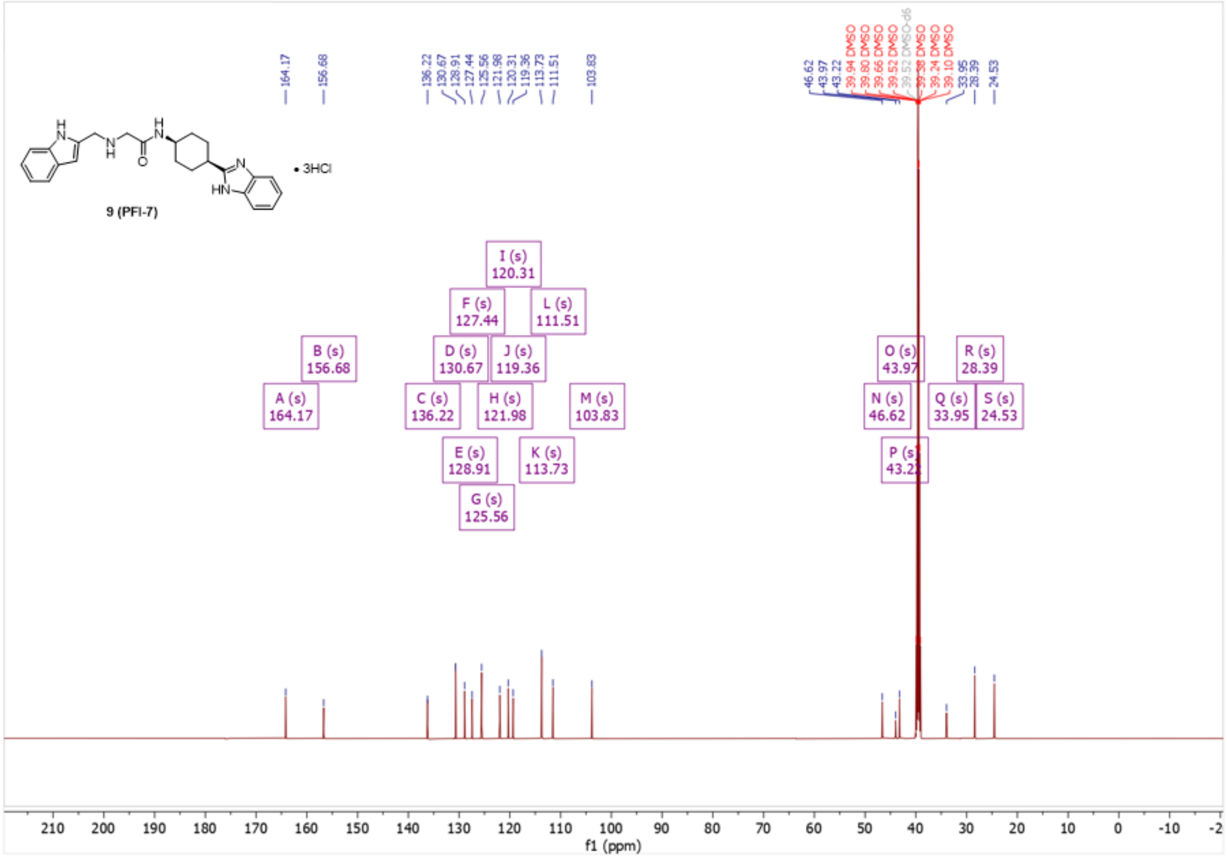

9 (PFI-7)
• 3HCl
A (s) 164.17
B (s) 156.68
C (s) 136.22
D (s) 130.67
E (s) 128.91
F (s) 127.44
G (s) 125.56
H (s) 121.98
I (s) 120.31
J (s) 119.36
K (s) 113.73
L (s) 111.51
M (s) 103.83
N (s) 46.62
O (s) 43.97
P (s) 43.22
Q (s) 33.95
R (s) 28.39
S (s) 24.53
f1 (ppm)

**Compound 24:**

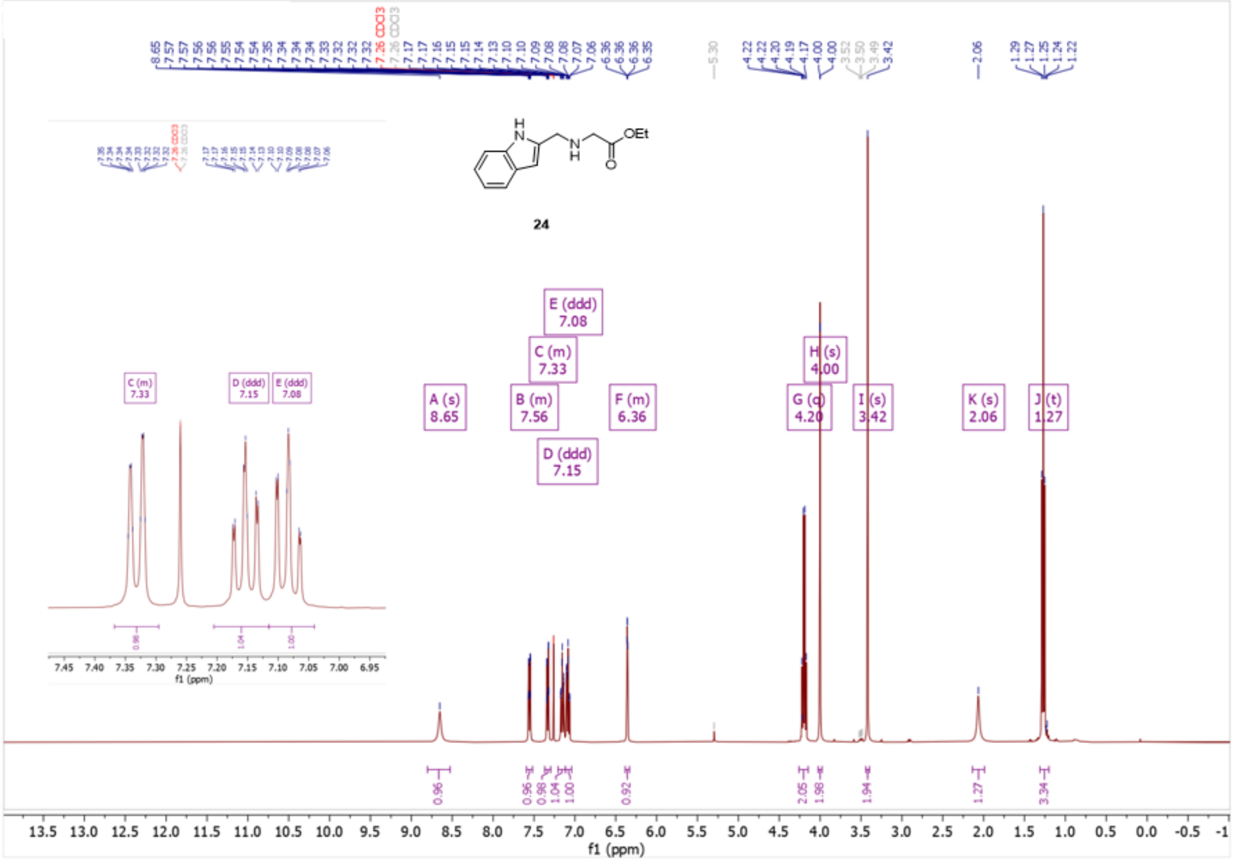

**Compound 25:**

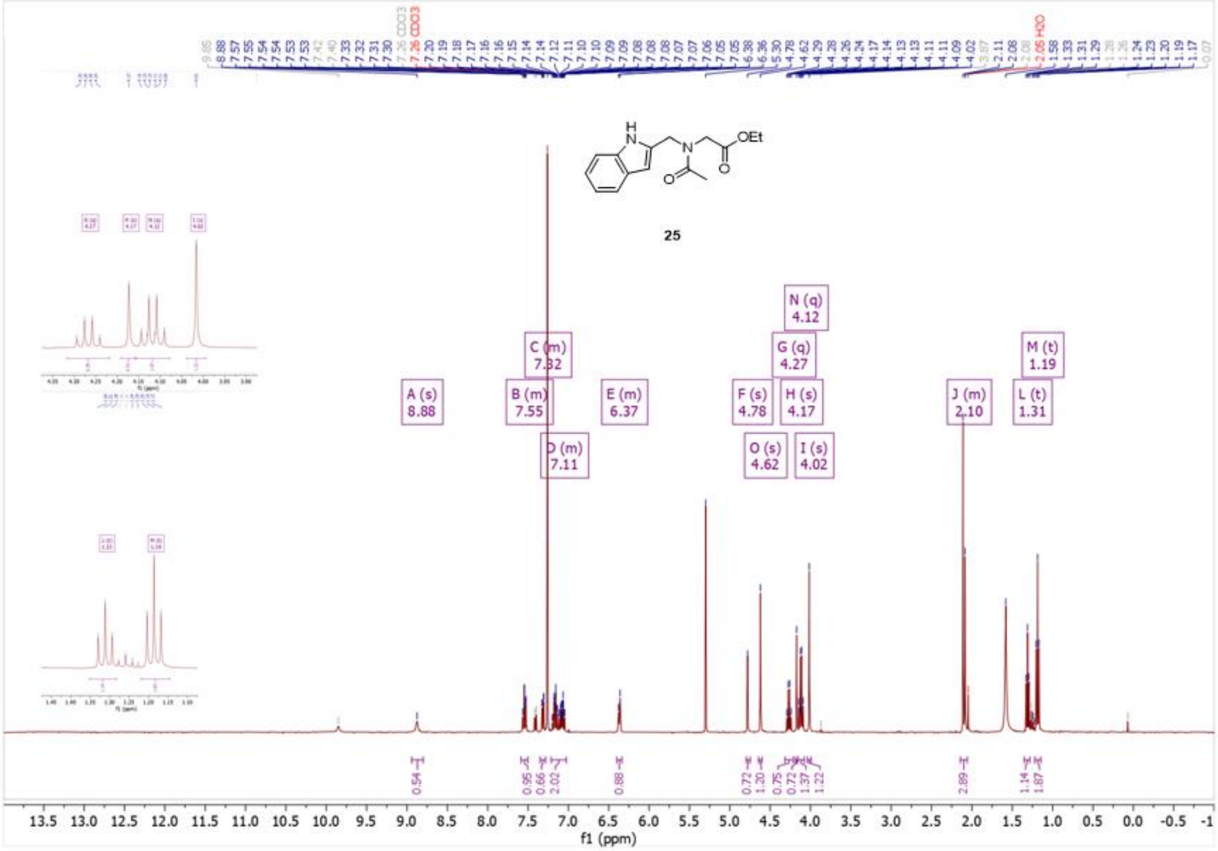

**Compound 26:**

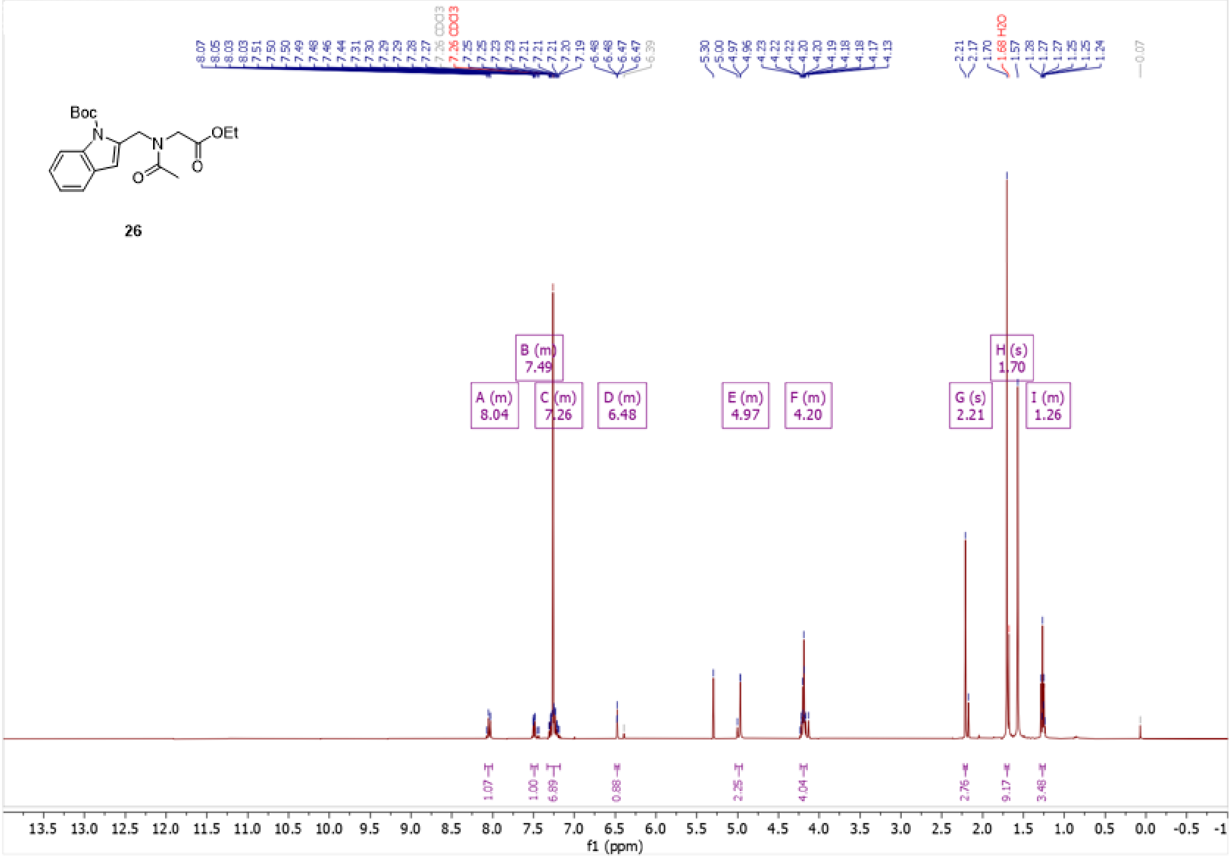

**Compound 27:**

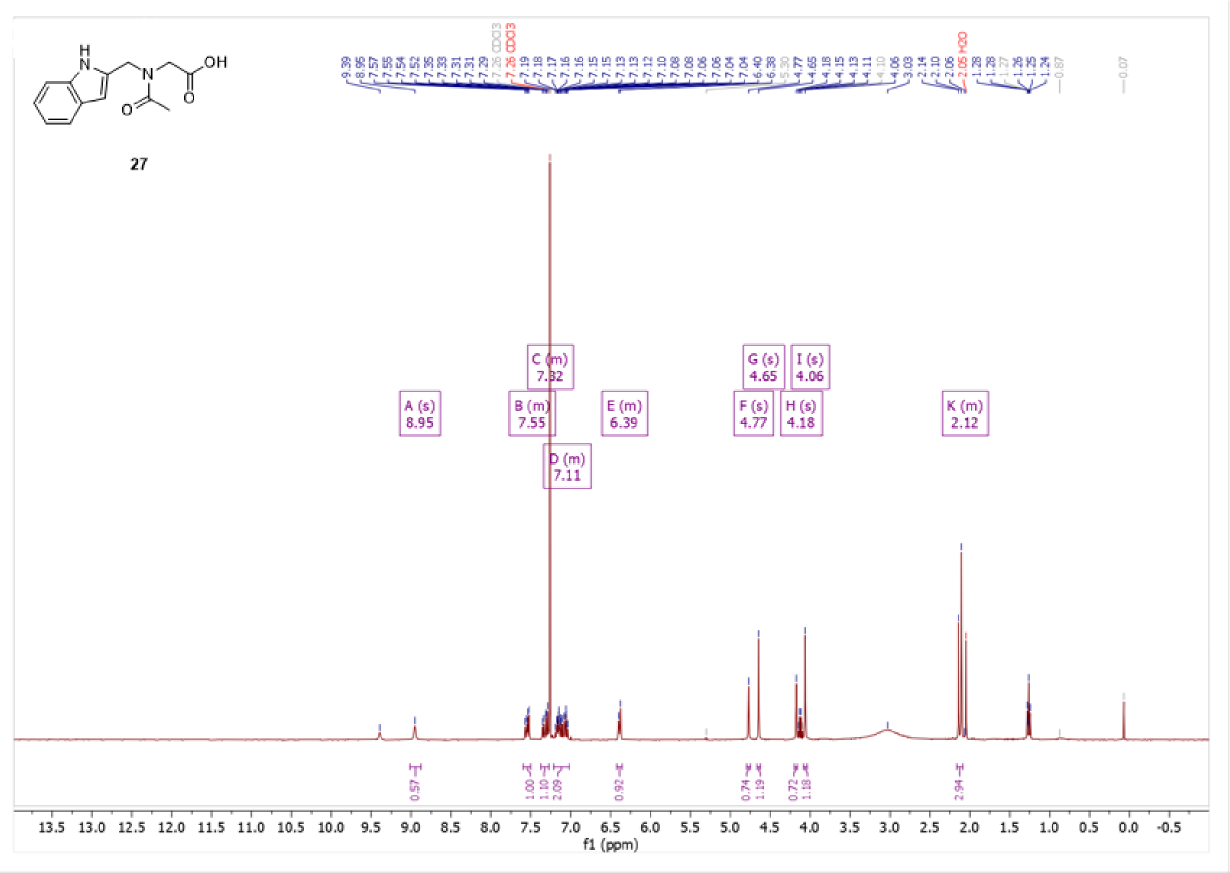

27
A (s) 8.95
B (m) 7.55
C (m) 7.32
D (m) 7.11
E (m) 6.39
F (s) 4.77
G (s) 4.65
H (s) 4.18
I (s) 4.06
K (m) 2.12
f1 (ppm)

**Compound 10, PFI-7N:**

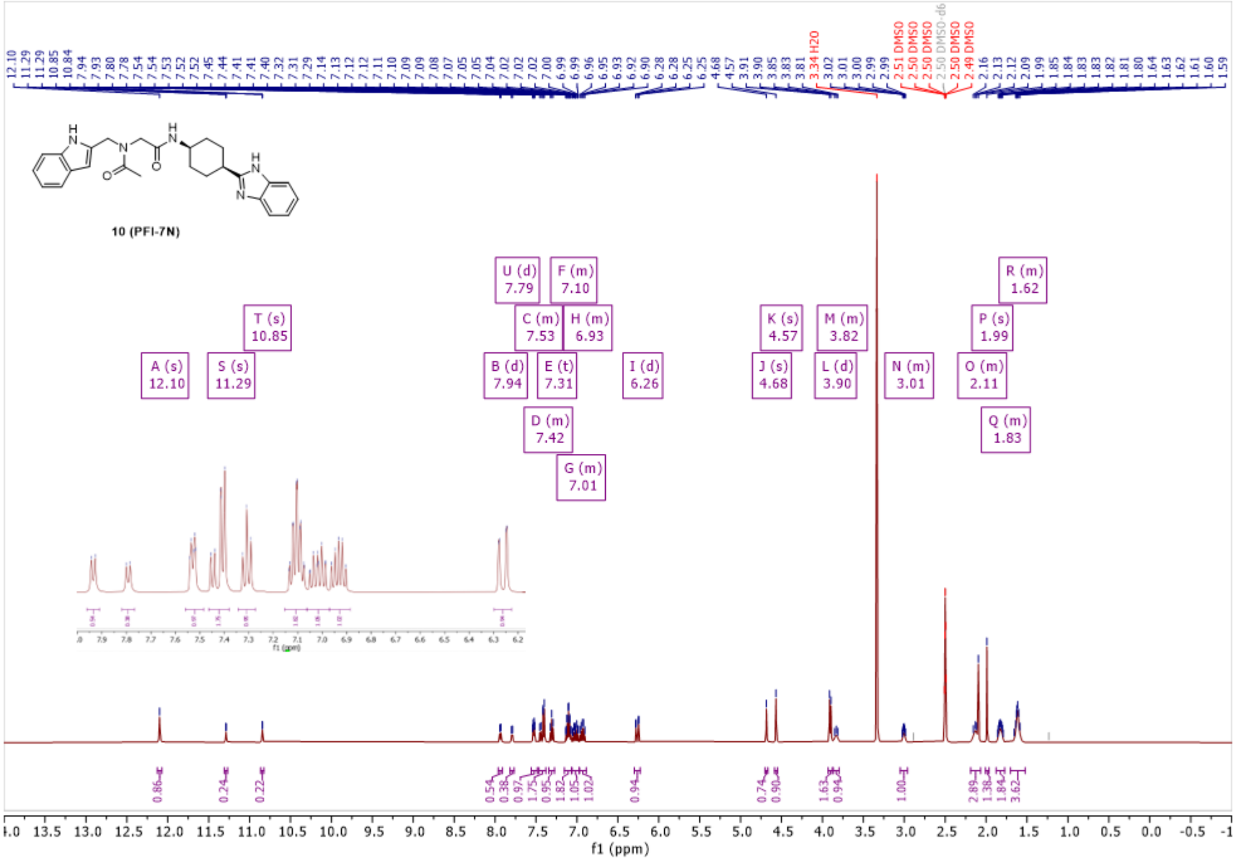

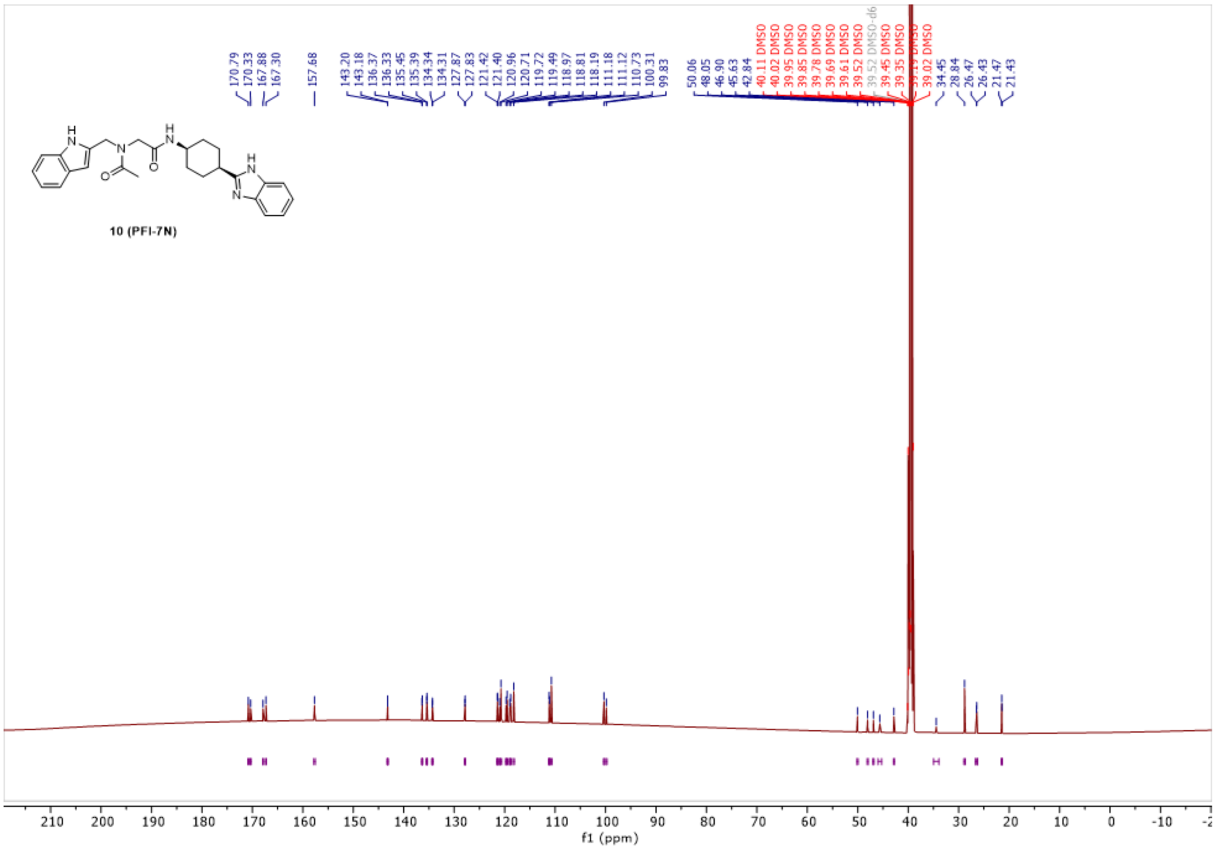

170.79
170.33
167.88
167.30
157.68
143.20
143.18
136.37
136.33
135.45
135.39
134.34
134.31
127.87
127.83
121.42
121.40
120.96
120.71
119.72
119.49
118.97
118.81
118.19
111.18
111.12
110.73
100.31
99.83
50.06
48.05
46.90
45.63
42.84
40.11 DMSO
40.02 DMSO
39.95 DMSO
39.85 DMSO
39.78 DMSO
39.69 DMSO
39.61 DMSO
39.52 DMSO
39.52 DMSO-d6
39.45 DMSO
39.35 DMSO
39.02 DMSO
34.45
28.84
26.47
26.43
21.47
21.43
H
N
H
N
O
O
H
N
N
10 (PFI-7N)
210 200 190 180 170 160 150 140 130 120 110 100 90 80 70 60 50 40 30 20 10 0 -10
f1 (ppm)

**Fluorescence polarization-based peptide displacement assay**

All Fluorescence Polarization (FP) experiments were performed in a total assay volume of 20 µL per well in 384-well black, low-volume, round-bottom polypropylene microplates (Greiner) using the C-terminally FITC labeled Pro/N-degron peptide (PGLWKS) and the GID4. For assessing the *K* displacement ($K_{disp}$) values, compounds were titrated in buffer (50 mM Tris(hydroxymethyl)aminomethane (Tris) buffer, pH 7.5, and 0.01% Triton X-100) for 2-fold dilution titration series (ranging from 0.009-150 µM for all compounds and incubated in a reaction mix containing 5 µM GID4, 40 nM C-terminally FITC-labelled PGLWKS and final DMSO concentration of 1.5% (v/v). FP was measured after 30 min of incubation at room temperature, using a BioTek Synergy 4 (BioTek, Winooski, VT) with excitation and emission wavelengths of 485 nm and 528 nm, respectively. The FP values were blank subtracted and were presented as the percentage of control (FP %) (Figure 1). All experiments were performed in triplicate (n = 3), and plotted values are the average of three replicates ± standard deviation. Data were visualized using GraphPad Prism software 7.04 (GraphPad, La Jolla, CA).

**Surface Plasmon Resonance (SPR)**

SPR studies were performed using a Biacore T200 (GE Health Sciences) at 25 °C. Biotinylated GID4 was captured onto a flow cell of a streptavidin-conjugated (SA) SPR chip (GE Healthcare) at approximately 5000 response units (RU) according to the manufacturer's protocol while another flow cell was left empty for reference subtraction. Serial dilutions of compounds (dilution factor of 0.33 was used to yield 10 concentrations) were prepared with 100 µM and for a few compounds (**5** and **7**) 10 µM as the highest compound concentration in HBS-EP+ Buffer (10 mM HEPES pH 7.4, 150 mM NaCl, 2 mM EDTA, 0.05% Tween 20, and 1 % DMSO). Kinetic determination experiments were performed using multi-cycle kinetics with 45 seconds (S) on time (i.e., association phase), and off time (disassociation time) of 60 (S) at a flow rate of 40 μL $min^{-1}$. HBS-EP+ Buffer only (plus 1% DMSO) was used for blank injections, and HBS-EP+ buffers containing 0.5 to 1.5% DMSO were used for buffer corrections. All experiments were performed in triplicate (n = 3). $K_D$ values were calculated by using steady-state affinity fitting and the Biacore T200 Evaluation software 3.1.

**NanoBRET PPI Assay**

HEK293T cells were seeded in 6-well plates (8e5 cells/well) and next day transfected with 0.2 µg/well C-terminally NanoLuc-tagged PGLWKS and 1.8 µg/well N-terminally HaloTag-tagged GID4 constructs, using Extreme gene HP transfection reagent, following manufacturer instructions. The following day cells were trypsinized and replated into 384-well white plates (2e5 cells/ml, 20 µl/well) in DMEM (no phenol red, 4% FBS) +/- 618 fluorescent ligand (1 µl /ml) and +/- indicated compound concentrations and DMSO. 4 h later the 5 µl Nanoglo substrate was added (8 µl/ml) to each well. Donor emission at 450 nm (filter, 450 nm/band-pass 80 nm) and acceptor emission at 618 nm (filter, 610 nm/long-pass) was measured within 10 min of substrate addition using a CLARIOstar microplate reader (Mandel). NanoBRET ratios were determined by subtracting 618/460 signal from cells without NanoBRET™ 618 Ligand x 1000 from 618/460 signal from cells with NanoBRET™ 618 Ligand x 1000. The IC50 was determined with GraphPad Prism software 7.04 (GraphPad, La Jolla, CA).

**Data Collection and Structure Determination:**

X-ray diffraction data for GID4 + UBF9092 was collected at 100K at beamline NSLS-II Beamline 17-ID-2 of Brookhaven National Laboratory. The data were processed using XDS (Kabsch, W. (2010a). XDS. Acta Cryst. D66, 125-132.) and the structure was solved by directly refined PDB entry 6WZZ as a model against the diffraction data with REFMAC (Murshudov GN, Vagin AA, Dodson EJ. Acta Crystallogr D Biol Crystallogr 1997;53:240–255) . REFMAC was also used for structure refinement. Graphics program COOT (Emsley P, Cowtan K. Acta Crystallogr D Biol Crystallogr 2004;60:2126–2132) was used for model building and visualization. Molprobity (Williams et al. (2018) MolProbity: More and better reference data for improved all-atom structure validation. Protein Science 27: 293-315.) was used for structure validation.

**Supplementary Table 1.** Crystallographic data and refinement statistics

| | GID1 + Compound **7** |
|---|---|
| **PDB Code** | 7S12 |
| **Data collection** | |
| Space group | $P4_12_12$ |
| Cell dimensions | |
| *a*, *b*, *c* (Å) | 40.1, 40.1, 201.0 |
| $\alpha$, $\beta$, $\gamma$ (°) | 90.0,90.0,90.0 |
| Resolution (Å) (highest resolution shell) | 40.20-2.15(2.21-2.15) |
| Unique reflections | 121329 |
| $R_{merge}$ | 5.3(138.2) |
| $I/\sigma I$ | 23.0(1.8) |
| Completeness(%) | 100.0(76.2) |
| Redundancy | 12.4(12.9) |
| CC(1/2) | 0.999(0.762) |
| **Refinement** | |
| Resolution (Å) | 39.00-2.15 |
| No. reflections (test set) | 8748(973) |
| $R_{work/}$ $R_{free}$ (%) | 22.1/27.5 |
| No. atoms | |
| Protein | 1296 |
| Compound | 23 |
| B-factors (Å$^2$) | |
| Protein | 68.9 |
| Compound | 61.6 |
| RMSD | |
| Bond lengths (Å) | 0.010 |

| | |
|---|---|
| Bond angles (º) | 1.825 |
| Ramachandran plot % residues | |
| Favored | 93.9 |
| Additional allowed | 6.1 |
| Generously allowed | 0 |
| Disallowed | 0 |
